# Neuroprotection by chronic administration of Fluoroethylnormemantine (FENM) in mouse models of Alzheimer’s disease

**DOI:** 10.1101/2024.10.31.621224

**Authors:** Allison Carles, Aline Freyssin, Sarra Guehairia, Thomas Reguero, Michel Vignes, Hélène Hirbec, Gilles Rubinstenn, Tangui Maurice

**Author notes:** Correspondence to: Tangui Maurice, PhD, INSERM UMR_S1198, Université de Montpellier, cc 105, Place Eugène Bataillon, 34095 Montpellier cedex 5, France. Tel.: +33/04 67 14 3291.

## Abstract

**Background:** Fluoroethylnormemantine (FENM), a new Memantine (MEM) derivative, prevented amyloid-β[25–35] peptide (Aβ_25–35_)-induced neurotoxicity in mice, a pharmacological model of Alzheimer’s disease (AD) with high predictive value for drug discovery. Here, as drug infusion is likely to better reflect drug bioavailability due to the interspecies pharmacokinetics variation, we analyzed the efficacy of FENM after chronic subcutaneous (SC) infusion, in comparison with IP injections in two AD mouse models, Aβ_25–35_ –injected mice and the transgenic APP /PSEN1^∂E9^ (APP/PS1) line.

**Methods:** In Aβ_25–35_-treated mice, FENM was infused at 0.03-0.3 mg/kg/day during one week after Aβ_25–35_ injection. For comparison, FENM and MEM were administered IP daily at 0.03-0.3 mg/kg. In 10-month-old APP/PS1 mice, FENM was administered during four weeks by daily IP injections at 0.3 mg/kg or chronic SC infusion at 0.1 mg/kg/day. Memory deficits, spatial working memory and recognition memory, were analysed. Markers of neuroinflammation, apoptosis, oxidative stress, and amyloid burden in APP/PS1 mice, were quantified. Markers of synaptic plasticity such as PSD-95 and GluN2A/B/D subunits expression in hippocampus homogenates or synaptosomes were quantified in Aβ_25–35_-treated mice and synaptic long-term potentiation (LTP) in hippocampal slices was analysed in APP/PS1 mice.

**Results:** Deficits in spontaneous alternation and object recognition in Aβ_25–35_ mice were prevented by infused FENM at all doses tested. Similar effects were observed with the daily FENM or MEM treatments. Animals infused with 0.1 mg/kg/day FENM showed prevention of Aβ_25–35_-induced neuroinflammation, oxidative stress and apoptosis. FENM infusion restored Aβ_25–35_-induced alterations in synaptosomal PSD-95, GluN2A and P-GluN2B levels. GluN2D levels were unchanged whatever the treatment. In APP/PS1 mice, FENM infused or administered IP alleviated spontaneous alternation deficits, neuroinflammation, increases in Aβ_1-40_/Aβ_1-42_ and hippocampal LTP alteration.

**Conclusion:** These data confirmed the neuroprotective potential of FENM in the pharmacological Aβ_25–35_ and transgenic APP/PS1 mouse models of AD, with a superiority to MEM, and showed that the drug can be efficiently infused chronically.

## Introduction

Alzheimer’s disease (AD) is the main form of age-related dementia and neurodegenerative pathology in the world population. It accounts for about 70% of the dementia cases and represents more than 50 million AD patients worldwide [1]. AD is characterized by progressive cognitive dysfunction, memory loss, and a drastic change in personality [for review, 2]. At the neuropathological level, main hallmarks of AD are the extracellular accumulation of aggregating amyloid-β proteins (Aβ), forming senile plaques, intracellular neurofibrillary tangles, composed of abnormally phosphorylated tau protein, and a massive neuroinflammation [for reviews, 2-4]. Although knowledge on the complex neuroanatomical and functional alterations occurring in AD has markedly increased these last decades, AD etiology is still controversial [5,6]. Many risk factors have been identified, such as oxidative stress, genetic mutations, endocrine disorders, age-related inflammation, depression, hypertension, infection, diabetes, food supplements, exposure to chemicals, metabolic conditions, and vitamin deficiencies [for review, 7]. While their connection to the susceptibility or the disease severity has been established, there is still no consensus hypothesis regarding AD pathogenesis, particularly in sporadic forms. The major roles of the pathological accumulation of amyloid species [3], tau hyperphosphorylation [8], neuroinflammation and oxidative stress [9] remain central in the progression of the neurodegeneration. It results in failure of neurons to maintain functional dendrites, synapse loss, and neuronal death affecting cholinergic neurons in several brain structures including the hippocampus and neocortex [10,11]. This neurodegeneration mainly originates from alteration of synaptic Ca^2+^ regulation in response to the over-activation of the ionotropic N-methyl-D-aspartate type of glutamate receptors (NMDARs) [12]. Physiological activity of NMDARs is necessary to mediate optimal synaptic transmission and normal neuronal function [13,14]. In the brain of AD patients, high levels of glutamate can be released from neuronal or glial cells, and reuptake, notably through astroglia, is altered, favoring massive influx of Ca^2+^ mediated by NMDAR and excitotoxicity [15]. The latter is responsible for generating oxidative stress and neuroinflammation and therefore promoting synapse loss and later, cell death [16,17].

Recent approvals of several passive immunotherapies targeting the pathological accumulation of amyloid species with monoclonal antibodies (Aducanumab, Lecanemab, Donanemab [18–20]) provided the first disease-modifying treatments in AD. However, these treatments carry risks and side effects, such as edema and hemorrhage, highlighting the ongoing need for effective pharmacological options in Alzheimer’s disease. Pharmacological treatments are based on acetylcholinesterase inhibitors, such as Donepezil, Rivastigmine, or Galantamine, to maintain the cholinergic tonus, and on Memantine (MEM), a NMDAR antagonist [12,21,22]. MEM acts as a non-competitive voltage-dependent NMDAR antagonist with moderate affinity, functioning as an open NMDAR channel blocker with fast off-rate, while also showing preferential blockade of extrasynaptic NMDARs [22–24]. MEM modulates the effects of pathologically elevated levels of glutamate that could lead to neuronal dysfunction and neurodegeneration.

In search of second-generation NMDAR biomarkers, several derivatives of MEM have been synthetized and tested as positron emission tomography radiotracers for the *in vivo* labeling of NMDARs [25]. [^18^F]-Fluoromemantine and [^18^F]-Fluoroethylnormemantine ([^18^F]-FENM) showed substantial brain accumulation and high *in vitro* affinity. However, unlike [^18^F]-Fluoromemantine, only [^18^F]-FENM demonstrated specific and selective binding profiles *in vivo* in rats, both under physiological conditions and in a brain injury model [25–29], binding to gray matter, *e.g.*, cerebral and cerebellar cortex and central grey nuclei. Depending on the area, brain-to-blood ratios varies from 5 (brain stem) to 8 (cortex) with a distribution matching that of the NMDAR GluN1 subunit. The non-radioactive isotopologue FENM showed significant pharmacological activities along with optimal pharmacokinetic and pharmacodynamic (PK/PD) characteristics. First, FENM acts as prophylactic and/or antidepressant against stress-induced maladaptive behavior in a rodent model of post-traumatic stress disorders [30,31]. Second, it revealed to be an effective neuroprotectant in a mouse model of AD [32]. In the latter study, the symptomatic and neuroprotective activities of FENM were analyzed in comparison with MEM, in the pharmacological model of AD induced by intracerebroventricular (ICV) injection of oligomerized Aβ_25-35_ peptide in mice [33]. After Aβ_25-35_ injection, mice rapidly developed neuroinflammation, oxidative stress, apoptosis, and learning deficits reminiscent of AD toxicity [34,36]. MEM and FENM showed symptomatic anti-amnesic effects in Aβ_25-35_-treated mice. Following repeated daily systemic injection, both compounds prevented the Aβ_25-35_-induced memory deficits, oxidative stress, inflammation, apoptosis and cell loss, in the hippocampus and cortex [32]. Interestingly, FENM effects were systematically more robust than that observed with MEM while the drug did not produce direct amnesic effect at higher doses, a classical effect of MEM suggesting its superiority as a neuroprotectant.

AD is a chronic pathology, requiring long-term treatment, and MEM-induced benefit for the patient, with the current dosage formulations as high doses, has been tempered due to adverse side-effects such as abdominal pain, nausea, vomiting or anorexia, poor adherence to the medication complicated by problems of dysphagia and memory loss, and variation of the drug plasma levels [37–40]. Furthermore, treatment discontinuation may result from severe medical complications such as asthenia, malaise, hepatotoxicity or kidney failure [41]. Novel drug delivery forms, such as transdermal infusion systems, represent interesting alternatives that enhance usability, dosage, and compatibility. Ideally, the drug delivery form should ensure a one-off administration, pharmacologically effective plasma concentrations of the compound over a prolonged period, preferably for at least 24 h or several days, leading to progressive achievement of the steady-state, continuous plasma concentration monitoring and easier management by caregivers. One of the challenges for animal translation of the MEM results is that its pharmacokinetics in human differs strongly from what is observed in murine models, the long half-life leading to higher and stable plasma concentration in human [42]. Even if FENM pharmacokinetics in human is not yet known, it is expected to be much longer than in mice, due to the standard interspecies animal variation. Moreover, infusion administration may also result in more translational pharmacodynamics in murine models than one-shot daily administration.

We here first compared the pharmacological effects of FENM in Aβ_25-35_-treated mice under two administration protocols during one week. Repeated daily intraperitoneal (IP) injections and continuous subcutaneous (SC) infusion using osmotic pumps. MEM, administered using repeated daily IP injections, was used as a control treatment. The drug-induced effects were analyzed in terms of memory alteration, using the spontaneous alternation and object recognition tests, neuroinflammation, using immunofluorescence and ELISA analyses oxidative stress, apoptosis markers, postsynaptic density 95 protein (PSD-95) and NMDAR subunit expressions. Changes in GluN2A/B/D subunits in the mouse hippocampus were investigated in homogenates and synaptosomal preparations. FENM was then tested in the APP_swe_/PSEN1^∂E9^ mouse line, using the optimal dosages identified in Aβ_25-35_-treated mice, namely: 0.3 mg/kg for repeated IP injection and 0.1 mg/kg/day for continuous SC infusion. Mice were treated at 10 months of age, when AD symptoms are established [43–46] and during four weeks. The drug-induced effects were analyzed in terms of memory alteration using the spontaneous alternation test, neuroinflammation, apoptosis markers, amyloid load, and analysis of long-term potentiation in hippocampal slices. The results confirmed the potentiality of FENM as a neuroprotectant in the two AD mouse models, the drug showing a complete protection when infused at a daily dosage as low as 0.1 mg/kg/day.

## Methods

### Animals

Male C57Bl/6j mice, from Janvier (Le Genest Saint Isle, France) were used at 5 weeks of age. Males and females APP_swe_/PSEN1^∂E9^ mice, co-expressing two genetic mutations associated with familial forms of AD, a humanized amyloid precursor protein (APP) carrying the Swedish (K670N_M671L) mutation (APP_swe_) and a mutant exon-9-deleted variant of the presenilin-1 (PSEN1^∂E9^) [43], thereafter referred to as APP/PS1 mice, bred on a C57BL6/J strain at the animal facility of IGF (French ministry of agriculture approval D34-172-13) from founders purchased at the Jackson laboratories (Bar Harbor, ME, USA) and subsequently transferred at 9-months old (mo) in the animal facility of the University of Montpellier (CECEMA, French ministry of agriculture approval E-34-172-23). Control littermates are thereafter referred to as wildtype (WT) mice. Mice were housed in groups of 9 mice maximum, except after the pump implantation surgery when they were housed individually, with free access to food and water and in a regulated environment (23°C ± 1°C, 40%–60% humidity, 12-hour-light/-dark cycle). Animal procedures were conducted in adherence with the European Union Directive 2010/63 and the ARRIVE guidelines [47] and authorized by the National Ethic Committee (Paris, France): authorization APAFIS #30410-2021031516372048.

### Drugs and peptides

FENM was from M2i Life Sciences (Saint Cloud, France). MEM was from Sigma-Aldrich (Saint Quentin-Fallavier, France). Drugs were solubilized in physiological saline (NaCl 0.9%, vehicle solution) as a stock solution (2 mg/ml corresponding to the dose of 10 mg/kg) and dilutions were done from this stock solution. The stock solutions were stored at +4°C up to 2 weeks. Drugs were administered using two administration protocols for 7 days: either repeated IP injections in a volume of 100 µl per 20 g body weight, or infusion using an osmotic micro-pump delivering 0.5 µl/h (see below). The amyloid-β[25–35] peptide (Aβ_25–35_) was from Genepep (St-Jean de Vedas, France). It was solubilized in distilled water at 3 mg/ml and stored at −20°C until use. Before injection, the peptide was incubated at 37°C for 4 days, allowing oligomerization. Control injection was performed with vehicle solution (distilled water) as we previously described no effect of antisense or control peptide, and ICV injections were done as described [33].

### Surgical procedures

In Aβ_25–35_ mice, osmotic micro-pumps model 1007D (Alzet, Charles River, France) were used with a capacity of 100 µl for infusion for 7 days. They were aseptically implanted at 6 weeks and infused at 0.5 µl/hour for 7 days. In APP/PS1 mice, osmotic micro-pumps model 1004 (Alzet, Charles River, France) were used with a capacity of 100 µl for infusion for 28 days. They were aseptically implanted at 10 months and infused SC at 0.11 µl/h for 28 days. The mice were briefly anesthetised with 2.5% isoflurane gas using a small nose cone attached to a modified bane circuit. The animal coat on the dorsal surface of the mice was trimmed and swabbed with betadine and lidocaine cream for local anaesthesia. A sterile blade was used to make a 1 cm incision perpendicularly to the base of the mice neck. Subcutaneous tissue was spread using a haemostat to create a pocket to pass a diameter of the pump. The pump was inserted into the pocket and the incision was closed with staple. On the incision the lidocaine cream was applied and the mouse was placed in the cage with heating until the recovery from the surgery. The entire procedure took an average of 10 min and did not exceed 15 min. Mice were monitored and weighed daily.

### Experimental series

In the Aβ_25-35_ mouse model, we examined the neuroprotection induced after two modes of administration of the drugs. First, drug infusion for 7 days was analysed in Aβ_25-35_-treated mice. Second, repeated administration by repeated daily IP injection for 1 week starting on the day of peptide injection was analysed. For the two modes of administration, two behavioural tests were performed: spontaneous alternation and object recognition test. Then, neuroprotection was analysed only for the FENM infused group. Animals were sacrificed at day 11, their hippocampus and cortex dissected out and flash frozen to assess cytokine levels by enzyme-linked immunosorbent assays (ELISA), homogenate or synaptosomal extraction to realize western blot and colorimetric assays. A second batch of mice was sacrificed by transcardiac paraformaldehyde fixation, to perform immunofluorescent analyses. In the APP/PS1 mouse line, three experimental groups were compared in 10-mo WT and APP/PS1 mice: (1) animals infused SC during 28 days with physiological saline (Veh); (2) animals infused SC during 28 days with FENM at the dose of 0.1 mg/kg/day; and (3) animals administered IP once-a-day (o.d.) during 28 days with FENM 0.3 mg/kg or MEM 0.3 mg/kg. Veh-infused WT and APP/PS1 mice served as controls also for the repeated IP injection of FENM condition, as it appeared to be the most drastic condition and controls were not duplicated to convey the 3R (reduce, refine, replace) rule in conducting animal experimental research [47]. The experimental group was divided in 2 batches. The first was euthanized by decapitation after anaesthesia for biochemical assays. The second was used for electrophysiological experiments.

### Behavioural Testing

Spontaneous alternation performance in the Y-maze, an index of spatial working memory, was measured as previously described [33–36]. Mice were allowed to explore the maze for 8 min. The series of arm entries, including possible returns into the same arm, was checked and the percentage of alternation was calculated as (actual alternations / number of arm entries – 2) x 100. Animals that performed less than 8 arm entries or showed alternation percentage < 20% or > 90% were discarded from the calculations. Recognition memory was analyzed using a novel object test [35,36]. Mice were placed individually in a squared open-field. In session 1, animals were allowed to acclimate during 10 min. In session 2, after 24 h, two identical objects were placed at ¼ and ¾ of one diagonal of the 50 x 50 cm^2^ arena. The mouse activity and nose position were recorded during 10 min (Nosetrack^®^ software, Viewpoint). The number of contacts with the objects and duration of contacts were measured. In session 3, after 24 h, the object in position 2 was replaced by a novel one differing in color, shape and texture. Each mouse activity was recorded during 10 min and analyzed. A preferential exploration index was calculated as the ratio of the number (or duration) of contacts with the object in position 2 over the total number (or duration) of contacts with the two objects. Animals showing no contact with one object or less than 10 contacts with objects, during the session 2 or 3, were discarded from the study. Animals did not receive drug treatment before the sessions.

### Brain fixation and slicing

After the behavioural tests, 30 min before the anaesthesia, buprenorphine (0.1 mg/kg) was injected SC for 5–6 mice from each condition. Mice were anesthetized with Exagon (1 ml/kg IP) and transcardially perfused with 50 ml of saline solution followed by 50 ml of Antigenfix (Diapath). The samples were kept for 48 h post fixation in Antigenfix solution at 4°C. Brains were immersed in a sucrose 20% Antigenfix solution and sliced within 1 month. Each brain was sliced in an area including the cortex, the *nucleus basalis magnocellularis*, and the hippocampal formation, between Bregma +1.80 to −2.80 according to Paxinos & Franklin [48]. Serial coronal frozen sections (25 μm thickness) were cut with a freezing microtome (Microm HM 450, ThermoFisher), collected in a 24-well plate, and stored in cryoprotectant at −20°C. Slices were placed on glass slides, each containing 6 coronal sections from 1 mouse.

### Lipid peroxidation

Mice were killed by decapitation, brains were rapidly removed, and the cortex dissected out, weighed, frozen in liquid nitrogen, and stored at –80°C until assayed. The level of lipid peroxidation was determined using the modified xylenol oxidation method as previously described [34,35].

### Enzyme-linked immunosorbent assays

Protein contents in tumour necrosis factor-α (TNFα), interleukin-6 (IL-6), B-cell lymphoma 2 (Bcl-2), Bcl-2–associated X (Bax), Aβ_1-40_ and Aβ_1-42_ were analysed using commercially available ELISA kits (see Supplementary Table 1). For n = 6 animals, one hippocampus was used. The tissue was homogenized after thawing with fresh lysis buffer (RIPA buffer with 3 detergents, pH 7.5), protease and phosphatases inhibitors were added (Complete mini, PhosSTOP, Roche Diagnostics, Meylan, France) and sonicated on ice for 4 × 5 s. After centrifugation (10,000 *g*, 5 min, 4°C), supernatants were then aliquoted and stocked at –80°C and used within 1 month for ELISA according to the manufacturer’s instructions. To analyse the level of Aβ_1-40_ and Aβ_1-42_ in the cortex of APP/PS1 mice, a guanidine extraction buffer (5 M guanidine HCl) and a centrifugation at 16,000 *g*, 20 min at 4°C were used to separate soluble (supernatant) and insoluble fraction (pellet). For each assay, absorbance was read at 450 nm and sample concentration was calculated using the standard curve. Results are expressed in pg of marker per mg of sample tissue.

### Immunohistochemical labelling of microglia (IBA-1) and astrocytes (GFAP)

For immunohistochemical labelling, slices in 24-well plates were incubated overnight at +4°C with Rabbit polyclonal anti-allograft inflammatory factor 1 (anti-IBA-1; 1:250; see Supplementary Table 2) and mouse monoclonal anti-glial fibrillary acidic protein (anti-GFAP; 1:400). Then, slices were incubated 2 h at room temperature with secondary anti-rabbit Cy3 cyanine dye (1:400) and secondary anti-mouse Alexa fluor 488 (1:1,000) antibodies. Slices were incubated 5 min with 4′,6-diamidino-2-phenylindole (DAPI) 10 μg/ml (1:50,000) and rinsed with potassium phosphate buffer saline. Finally, slices were mounted with Dako Fluorescence Mounting Medium (Dako). Images of each slice were taken with a fluorescent Microscope (Zeiss). Data from GFAP immunolabeling were calculated as average of 5-6 slices per animal, with 5 animals per group, and expressed as number of immunoreactive cells per mm^2^. Internal variability of the counting was 5.1 ± 0.4% in Rad, 4.8 ± 0.4% in Mol, and 4.0 ± 0.3% in PoDG. Data from Iba1 immunolabeling were calculated as average of 5 slices per animal, with 5 animals per group, and expressed as number of immunoreactive cells per mm^2^. Internal variability of the counting was 5.4 ± 0.4% in Rad, 5.9 ± 0.5% in Mol and 7.6 ± 0.6% in PoDG.

### Western blotting

Mice were killed at indicated days after injections and the hippocampus rapidly dissected on ice and kept at –80°C until use. To compare the quantities of antibodies in the hippocampus, two types of brain extraction are used. First, hippocampus was sonicated on ice for 4 × 5 sec with 300 µL of fresh lysis buffer (RIPA buffer with 3 detergents), protease and phosphatases inhibitors were added (Complete mini, PhosSTOP, Roche Diagnostics, Meylan, France). Homogenates were centrifuged at 14 000 rpm for 15 min at 4°C and the supernatants were then aliquoted and stocked at –20°C. Second, synaptosomal hippocampus extraction was realized. Hippocampi were dissociated mechanically on ice using a mini-potter in a solution of Syn-PER (Thermo, 87793) containing protease and phosphatases inhibitors were added (Complete mini, PhosSTOP) at the concentration of 10 mL/g tissue. The samples were then centrifuged at 1,200 *g*, 4°C for 10 min. The supernatant was separated and a part centrifuged at 15,000 *g*, 4°C for 20 min. The supernatant from the second centrifugation corresponded to the cytosolic fraction and was stored at –20◦C. The pellet, corresponding to the synaptosome fraction, was resuspended with Syn-PER+PI at a concentration of 1 mL/g of initial tissue. Proteins contents after both extractions were quantified with BCA protein assay kit (Pierce Thermo-Scientific). Total proteins (30 μg) were diluted in sodium dodecyl sulphate loading buffer. Samples were loaded in 7.5% polyacrylamide resolving gel and electroblotted into a polyvinylidene fluoride membrane (Immobilon Transfer membrane, IPVH00010, Merck Millipore, Cork, Irlande). Blots were blocked in phosphate buffer saline (PBS) containing 0.1% or 0,05% Tween-20 (PBS-T) and 5% non-fat dry milk for 1 h and incubated with primary antibodies overnight at 4°C for anti-postsynaptic density 95 mouse antibody (anti-PSD-95 [6G6-1C9]; 1:1000; see Supplementary Table 2) and anti-nitrotyrosine mouse antibody (1:1000; Supplementary Table 2). For other antibodies, blots were blocked in Tris-buffered saline (TBS) containing 0.1% Tween-20 (TBS-T) and 5% bovine serum albumin for 2 h and incubated with primary antibodies overnight at 4°C. Primary antibodies used are detailed in Supplementary Table 2. After 3 washes of 5 min with PBS-T, membranes were incubated 1 h with horseradish peroxidase (HRP)-conjugated anti-rabbit antibody or anti-mouse antibody (each 1:2000; Supplementary Table 2), then washed 3 times for 10 min. To visualize the signal, blots were incubated in Immobilon^®^ crescendo western HRP substrate (WBLUR0500; Millipore) during 1 min. Pictures of the chemiluminescence signals were taken with Chemidoc System (Bio-Rad). Normalization was performed with the total protein using Stain-Free™ protocol. Stain-free technology eliminated many of the issues related to signal normalization for accurate protein quantification. Total protein level was directly measured on the gel used for western blotting. The total density for each lane was measured without need to strip and reprobe blots for housekeeping proteins to normalize protein levels, using the total lane profile instead [49,50]. Each blot was striped by solution Reblot plus strong 1X (10X, Millipore, 2504) during 15 min. Blot was blocked in PBS-T 0.1% or 0,05% and 5% non-fat dry milk for 1 h or Tris buffer saline-T 0.1% and 5% bovine serum albumin for 2 h and incubated with primary antibodies overnight at 4°C.

### Electrophysiological analyses of LTP

Experiments were carried out on freshly prepared hippocampal slices (250 μm) obtained from 11-m.o. APP/PS1 mice. After decapitation, the brains were quickly dissected and placed in ice-cold cutting buffer containing 40 mM NaCl, 4 mM KCl, 26 mM NaHCO_3_, 1.25 mM NaH_2_PO_4_, H_2_0, 0.5 mM CaCl_2_, 2 H_2_0, 7 mM MgCl_2_, 6 H_2_0, 10 mM glucose and 150 mM sucrose at pH 7.4 and 330 mOsmol (infused with O_2_/CO_2_, 95%/5%). The slices were then prepared using a vibratome (Leica VT 1000S) and kept at 25°C for at least 1 h before electrophysiological recording in artificial cerebrospinal fluid (125 mM NaCl, 3.5 mM KCl, 26 mM NaHCO_3_, 1.2 mM NaH_2_PO_4_, H_2_0, 2.4 mM CaCl_2_, 2 H_2_0, 1.3 mM MgSO_4_, 6 H_2_0 and 25 mM glucose) at 7.35 and 310 mOsmol. This latter buffer was used for the recordings. For electrophysiological recordings, the slices were transferred to a multielectrode array (MEA 2100-Mini, Multi Channel Systems, Reutlingen, Germany) continuously perfused with the extracellular medium described above (flow rate 1-2 ml/min) and maintained at 32°C. The MEA consisted of 64 extracellular electrodes. The inter-electrode distance was 200 μm. Each individual electrode in the array can be used for recording or stimulation. A nylon mesh was positioned over the slice to achieve satisfactory electrical contact between the surface of the slice and the electrode array. Biphasic stimulation (–100 µA to 100 µA) was delivered by one stimulator (MCS-SCU in vitro, IFB-C, Multi Channel Systems) in the Schaffer collateral pathway and recordings were made in the CA1 area of the hippocampus [51]. Basal neurotransmission was elicited by stimulations spaced 15 s apart. In order to achieve a long-term potentiation (LTP) process, a high frequency stimulation (HFS) consisting in 100 Hz pulse for 1 s was applied to CA1 afferents. Postsynaptic field excitation potentials (fEPSP) were then recorded by all remaining electrodes in the array at the same time. The signals were recorded and analyzed (MC Analyzer, Multi Channel Systems).

### Statistical analyses

Analyses were done using Prism v9.1 (GraphPad Software, San Diego, CA, USA). Data were analyzed using one-way analyses of variance (ANOVA, *F* value) or using two-way ANOVA with the ICV or genotype and drug treatments as independent factors, followed by a Dunnett’s *post-hoc* test. Biochemical analyses (Western blotting, Elisa, colorimetric assays) were analyzed using a Kruskal-Wallis non-parametric ANOVA followed by a Dunn’s *post-hoc* test, as the number of samples per group was low and/or groups showed heterogeneity of variances. Object preference, calculated from the duration of contacts with the two objects, was analyzed using a one-sample *t*-test *vs*. the chance level (50%). Significance levels were *p* < 0.05, *p* < 0.01 and *p* < 0.001. Statistical data are all reported in the Supplementary Table 3.

## Results

### FENM treatment by SC infusion or repeated IP injections improved memory impairment in A**β**_25-35_-treated mice

FENM was infused SC during 7 days in C57Bl/6 mice using an implanted osmotic micropump, in the 0.03 to 0.3 mg/kg/day dose-range. Infusion started on day 1, when Aβ_25-35_ was acutely administered ICV until day 7, *i.e.*, 24 h before the start of the behavioral analyses (Fig. 1a). This mode of administration was compared to repeated daily IP injections of FENM or MEM, in the same dose-range and as previously described in Swiss OF-1 mice [32].

**Figure 1.**
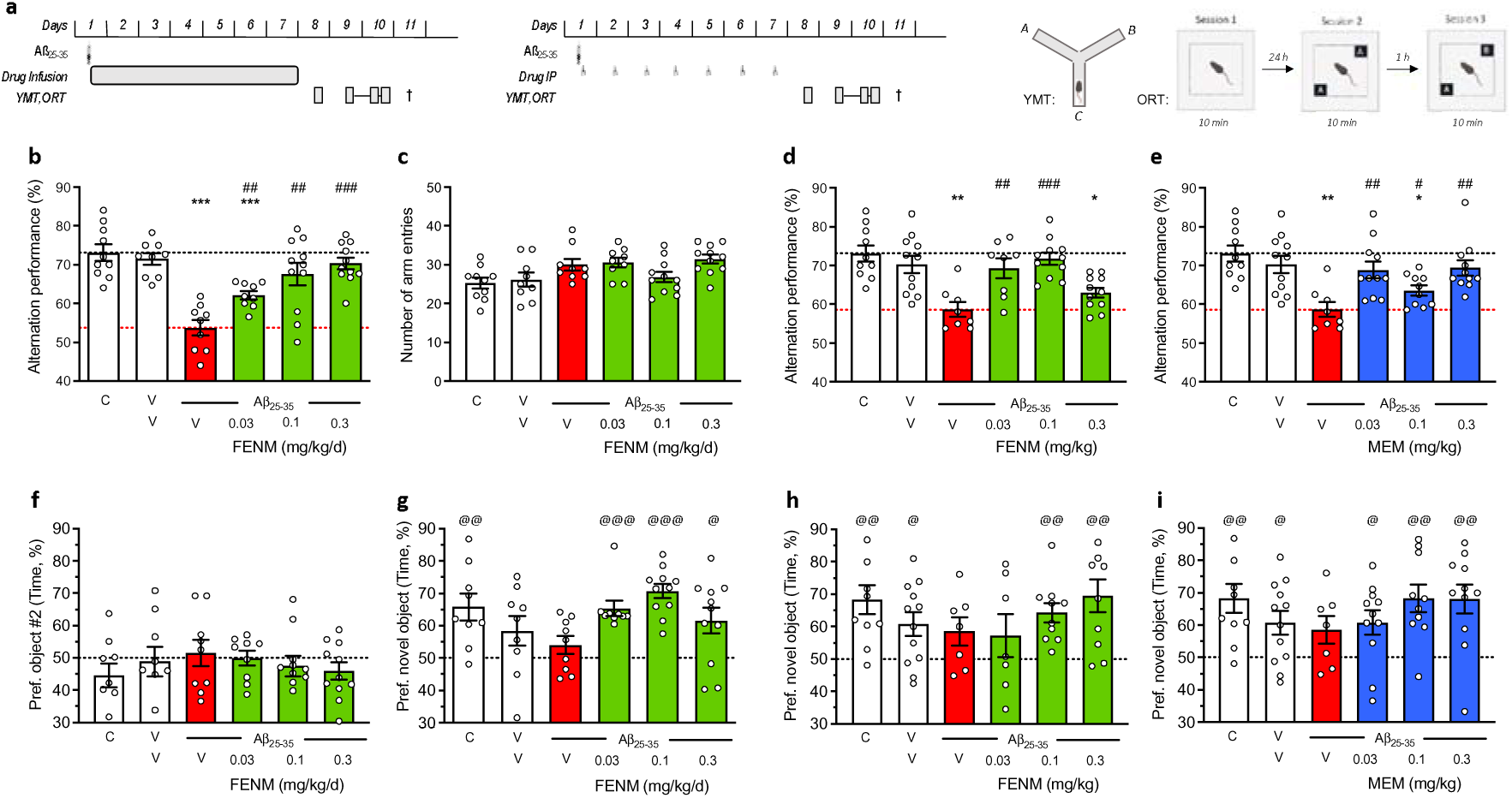
Protective effect of SC-infused Fluoroethylnormemantine (FENM) or IP-administered FENM or Memantine (MEM) on Aβ_25-35_-induced memory impairments in mice. (**a**) Experimental protocols for the SC drug infusion or drug IP administration. Abbreviations: YMT, Y-maze test; ORT, object recognition test; †, sacrifice before immunofluorescence or biochemical analyses. Spontaneous alternation in the Y-maze consists in recording the successive arm exploration in the maze over 8 min. The object recognition test consists in three 10-min duration sessions (i) without objects, (ii) with two identical objects, and (iii) with one familiar and one novel object. The object exploration preference was determined on the duration of contacts with each object. Animals received Aβ_25-35_ (9 nmol ICV) on day 1 and then FENM (0.03-0.3 mg/kg/day in the Alzet pump) or FENM or MEM (0.03-0.3 mg/kg IP o.d.) between day 1 to 7. (**b**, **d**, **e**) spontaneous alternation deficits and (**c**) number of arm entries. (**f**) session 2 of the object recognition test with two identical objects and (**g**, **h**, **i**) session 3 with a novel and a familiar object. Exploration preferences are calculated with the duration of contacts. Abbreviations: C, control (untreated) animals; V, vehicle solution (distilled water for ICV and physiological saline for IP injections of SC infusion. Graphs show individual data and mean ± SEM of N = 7-12 animals per group. Dotted lines show the (V+V)– and (V+ Aβ_25-35_)-treated group levels in (b-e) and the 50% preference level in (f-i). Abbreviations: C, control, non-injected animals; V, vehicle solution. * *p* < 0.05, ** *p* < 0.01, *** p < 0.001 *vs*. (V+V)-treated group; ## *p* < 0.01, ### *p* < 0.001 *vs*. (V+Aβ_25-35_)-treated group; Dunnett’s test. ^@^ *p* < 0.05, ^@@^ *p* < 0.01, ^@@@^ *p* < 0.001 *vs*. 50% level, one-sample *t*-test.

On day 8, mice were tested for their ability to spontaneously alternate when exploring the Y-maze, a measure of spatial working memory (Fig. 1b-e). Aβ_25-35_-treated mice receiving vehicle solution in the pump showed a highly significant deficit as compared with V/V-treated animals or untreated controls (Fig. 1b). The FENM infusion led to a dose-dependent attenuation of the Aβ_25-35_-induced deficit, with significant effects observed for all groups and complete recovery at 0.1 and 0.3 mg/kg/day (Fig. 1b). The treatments had no effect on locomotor and exploratory activities, as the number of arms explored during the 8-min session was unchanged among groups (Fig. 1c). For comparison, FENM administered repeatedly o.d. attenuated the Aβ_25-35_-induced alternation deficits at the doses of 0.03 and 0.1 mg/kg IP (Fig. 1d). MEM, administered IP, significantly attenuated the Aβ_25-35_-induced alternation deficit at all doses tested (Fig. 1e).

On days 9 and 10, animals were tested in a novel object test to analyze their recognition memory. The test consisted in a first session, on day 9, without object to train animals in the open-field. On day 10, two similar objects were presented and the interactions of each mouse with the object was analyzed. During this session 2, all groups receiving Aβ_25-35_ and/or FENM infusions showed no exploratory preference among the objects (Fig. 1f).

Session 3 was performed after 1 h and a familiar object was replaced but a novel one. Control mice showed a significant preferential exploration of the novel object that was not the case of Aβ_25-35_-treated animals (Fig. 1g). FENM infusion restored a preferential exploration of the novel object, with significant effects measured at all doses tested (Fig. 1g).

When administered repeatedly IP, both FENM (Fig. 1h) and MEM (Fig. 1i) significantly improved the novel object recognition in Aβ_25-35_-treated mice particularly at the doses of 0.1 and 0.3 mg/kg.

### FENM treatment by SC infusion or repeated IP injections improved A**β**_25-35_-induced toxicity

The anti-inflammatory activity of FENM infusion was first analyzed using global analyses of cytokine levels in the hippocampal tissue (Fig. 2a). TNFα level was significantly increased by Aβ_25-35_ treatment (+81%; Fig. 2b) and this increase was attenuated by FENM infusion, at all doses tested. IL-6 level showed a +37% (*p* = 0.126) trend to be increased by Aβ_25-35_ and FENM treatment, particularly at the 0.1 mg/kg/day dose, prevented this increase, but the ANOVA did not reach significance (Fig. 2c). Brains of FENM-infused mice were also analyzed for neuroinflammation using an immunofluorescence approach to visualize in the hippocampal formation both astroglial reaction, using GFAP expression (Fig. 2d-i), and microglial reaction, using IBA-1 expression (Fig. 2j-o). Three areas were examined: the *stratum radiatum*, the *stratum lacunosum-moleculare* and the polymorphic layer of the dentate gyrus. The Aβ_25-35_ treatment increased highly significantly GFAP immunolabelling in all three areas (Fig. 2d, f, h) by +29% to +87% (Fig. 2e, g, i) and the FENM infusion, at the dose of 0.1 mg/kg/day, significantly prevented these increases without altering GFAP expression alone. Similarly, the Aβ_25-35_ treatment highly significantly increased IBA-1 immunostaining in all three areas (Fig. 2j, l, n) by +45% to +53% (Fig. 2k, m, o). The FENM infusion, at 0.1 mg/kg/day, significantly prevented these increases without altering IBA-1 expression alone. These observations however convergently indicated that the SC infusion of FENM efficiently prevented the Aβ_25-35_-induced neuroinflammation in the mouse hippocampus.

**Figure 2.**
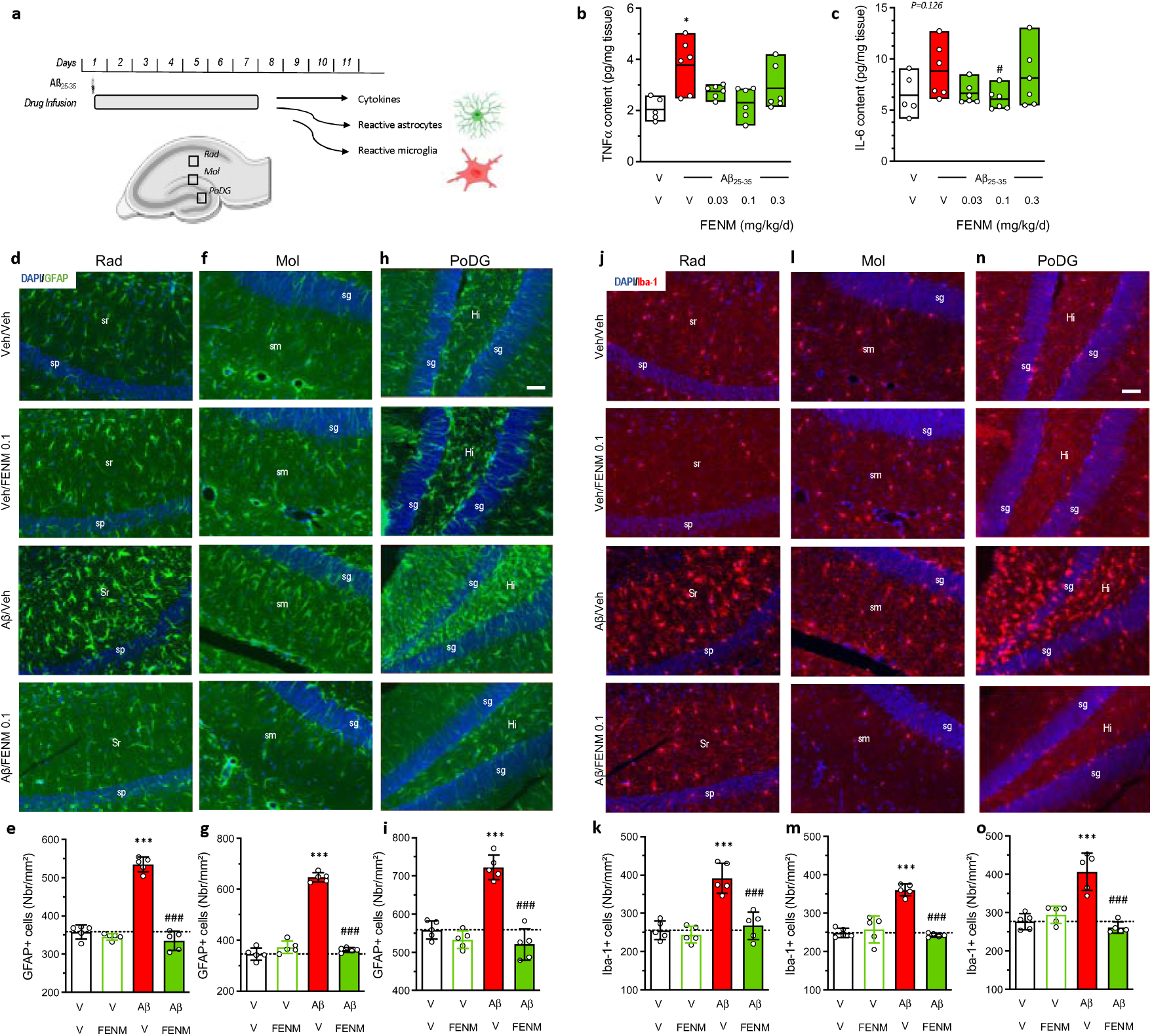
Protective effects of SC-infused FENM, 0.1 mg/kg/day, on Aβ_25-35_-induced neuroinflammation in mice. (**a**) Experimental protocol and hippocampal areas of interest. (**b**) TNFα and (**c**) IL-6 levels were analyzed in homogenates by ELISA. (**d**-**i**) Astroglial reaction was analyzed in the hippocampus of Aβ_25-35_-treated mice using GFAP immunolabeling. (**j**-**o**) Microglial reaction in the hippocampus of Aβ_25-35_-treated mice using Iba-1 immunolabeling. (**d**, **e**, **j**, **k**) *stratum radiatum* (Rad), (**f**, **g**, **l**, **m**) *stratum lacunosum-molec*ulare (Mol) and (**h**, **i**, **n**, **o**) polymorphic layer of the dentate gyrus (PoDG) with (**d**, **f**, **h**, **j**, **l**, **n**) typical immunofluorescence micrographs (blue: DAPI, green: GFAP, red: Iba-1) and (**e**, **g**, **i**, **k**, **m**, **o**) quantifications. Coronal 25 µm thick sections were stained with anti-GFAP or anti-Iba-1 antibody and the three areas of the hippocampus analyzed. Abbreviations: sp, *stratum pyramidale*; sr, *stratum radiatum*; sg, *stratum granulare*; sm, *stratum lacunosum-moleculare*; Hi, *hilus*. Scale bar in (h, n) = 50 µm applying to all pictures. The number of slice analyzed per mouse was n = 5-6 with N = 5 mice per group. *** *p* < 0.001 *vs*. the (V+V)-treated group; ### *p* < 0.001 *vs*. the (Aβ_25-35_+V)-treated group; Dunnett’s test.

Two parameters measuring Aβ_25-35_-induced oxidative stress was analyzed: the level of lipid peroxidation of cell membranes in the cortex (Fig. 3a) and the nitrosylated proteins in the hippocampus (Fig. 3b). Aβ_25-35_ significantly increased lipid peroxidation (+84%; Fig. 3a) and nitrosylated proteins (+35%; Fig. 3b). The FENM infusion significantly attenuated this increase at all doses tested (Fig. 3a, b).

**Figure 3.**
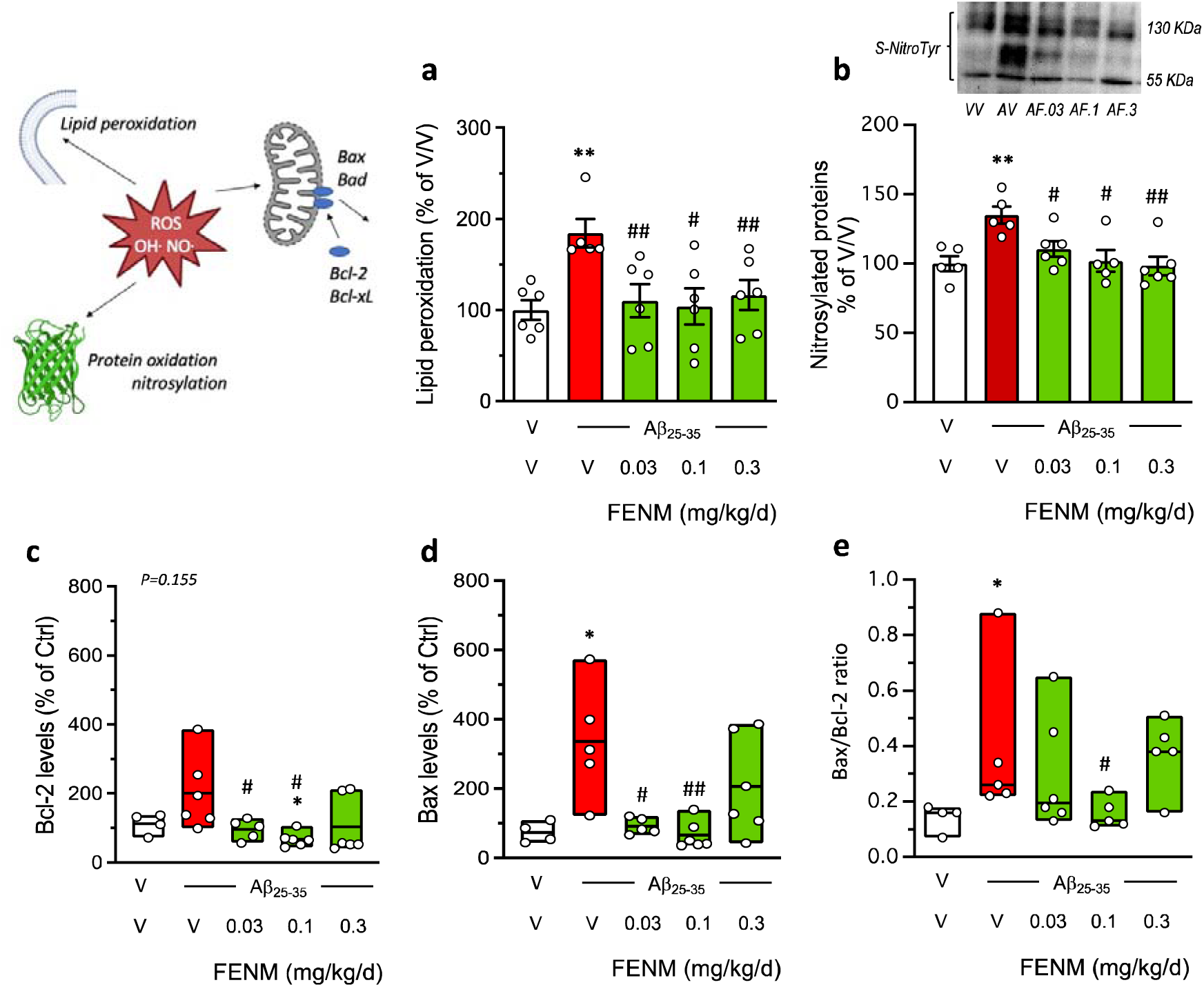
Protective effects of SC infusion of FENM, on Aβ_25-35_-inducedoxidative stress and apoptotic markers. (**a**) lipid peroxidation, (**b**) nitrated proteins, (**c**) Bcl-2 level, (**d**) Bax level, and (**e**) Bax/Bcl-2 ratio, measured by colorimetric assay, western blotting or ELISA in mouse hippocampus extracts. Animals were sacrificed 11 days after Alzet pump implantation. In (b), data show mean ± SEM and a typical blot is shown above the graph. In (c-e), data show median and min/max with individual measures. The number of animals per group was N = 4-6. * *p* < 0.05, ** *p* < 0.01 *vs*. (V+V)-treated group; # *p* < 0.05, ## *p* < 0.01 *vs*. (V+Aβ_25-35_)-treated group; Dunn’s test.

Aβ_25-35_ treatment induces apoptosis and cell death in the mouse hippocampus. We analyzed two proteases, Bcl-2 and Bax, known as cellular anti-apoptotic and pro-apoptotic markers, respectively. Both Bcl-2 and Bax levels were increased after Aβ_25-35_ treatment (+100% and +236%, respectively; Fig. 3c, d). Bax level was significantly increased by Aβ_25-35_, but not Bcl-2 (*p* = 0.155) (Fig. 3d). The FENM infusion attenuated these increases particularly at the 0.03 and 0.1 mg/kg/day doses. Moreover, the Bax/Bcl-2 ratio, considered as a reliable index of apoptosis level, was increased by Aβ_25-35_ (+170%; Fig. 3e) and this increase was attenuated in a U-shaped manner following infusion of FENM with a maximum effect at 0.1 mg/kg/day (Fig. 3e).

### FENM treatment by SC infusion improved A**β**_25-35_-induced NMDAR subunit altered expressions

Finally, in the Aβ_25-35_ model, we examined the impact of the FENM infusion on the expression of NMDAR subunits and the postsynaptic scaffolding protein PSD-95. First, we determined the time-course of protein expressions after Aβ_25-35_ ICV injection in hippocampus homogenates and synaptosomal preparations (Fig. 4). In homogenate preparations, the western blot analysis of PSD-95 expression showed a significant reduction at day 5 after the Aβ_25-35_ ICV injection, but not after 7 days (Fig. 4a). Neither GluN2A (Fig. 4b) nor GluN2B levels (Fig. 4c), and *a fortiori* the GluN2A/GluN2B ratio (Fig. 4d), were affected at days 5 or 7 after the peptide injection. In order the refine the analysis, we prepared synaptosomal preparations of the hippocampus. In synaptosomes, PSD-95 levels were significantly decreased at day 5 and 7 after Aβ_25-35_ injection (Fig. 4e). GluN2A levels were increased at day 5 and 7 after Aβ_25-35_ injection (Fig. 4f). GluN2B levels were unaltered (Fig. 4g), but the GluN2A/GluN2B ratio was significantly increased at both timepoints (Fig. 4h).

**Figure 4.**
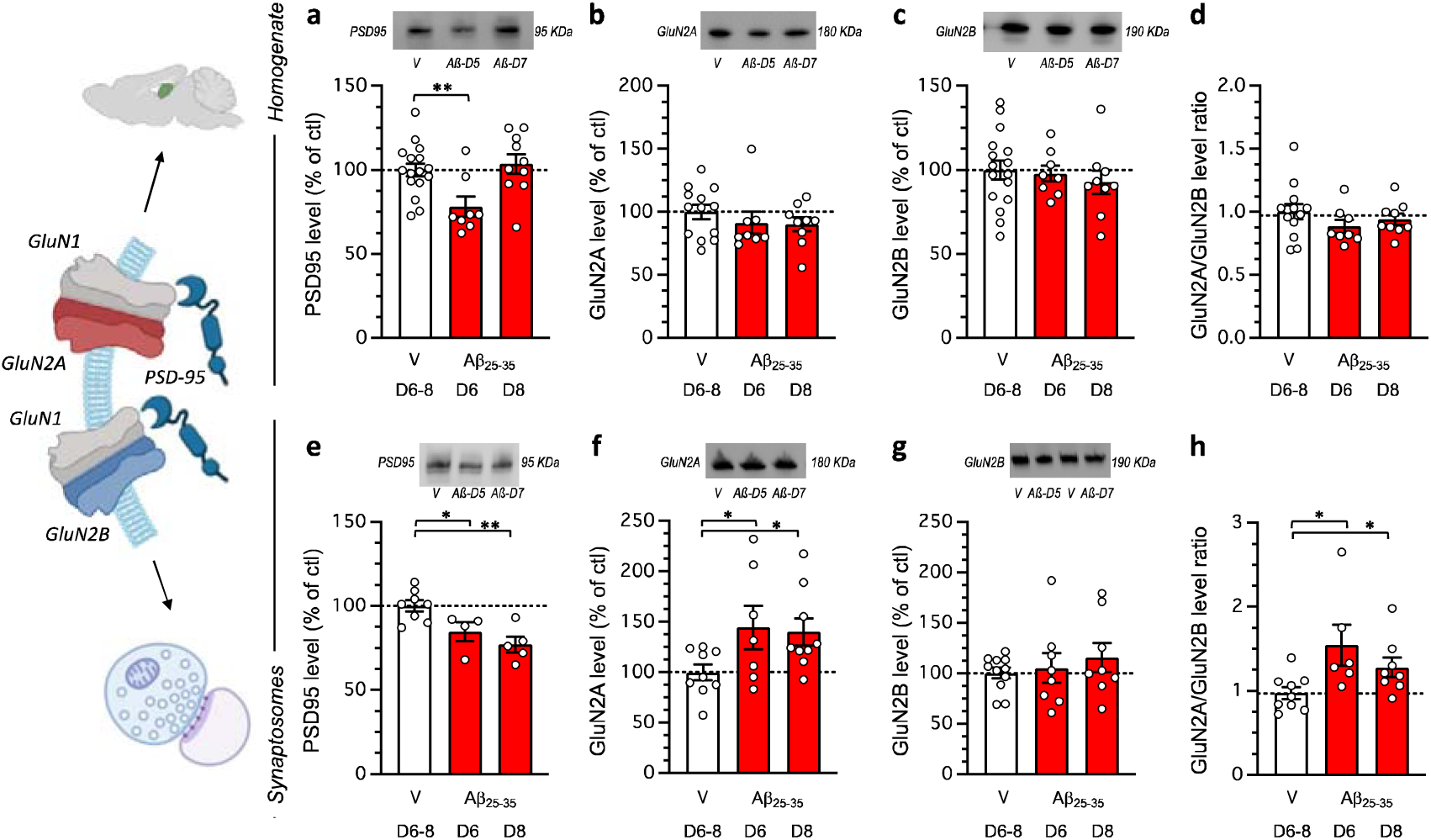
Impact of Aβ_25-35_ toxicity on PSD-95 and NMDAR GluN2A/B subunits in the mouse hippocampus. Aβ_25-35_ was administered ICV on day 1 and after 5 or 7 days, the hippocampus was dissected out and protein expression analyzed in homogenates. (**a**, **b**, **c**, **d**) or synaptosomes preparations (**e**, **f**, **g**, **h**) by western blot: PSD-95 levels (**a**, **e**), GluN2A expression (**b**, **f**), GluN2B expression (**c**, **g**) and GluN2A/GluN2B ratios (**d**, **h**). Note that protein levels were not altered on days 1 or 3 after Aβ_25-35_ peptide ICV injection (data not shown). Vehicle solution (V)-treated animals were sacrificed at day 6 or 8 and data were pooled. Typical blots are shown above each graph. The whole immunostained and and stain-free blots are shown in Supplementary Figures 1 and 2. The number of animals per group was N = 8-16 per group in (a-d) and N = 4-11 in (e-h). * *p* < 0.05, ** *p* < 0.01; Dunnett’s test.

The impact of the FENM infusion was therefore analyzed in synaptosomal preparations 7 days after Aβ_25-35_ ICV injection (Fig. 5). The drug, at 0.1 mg/kg/day restored PSD-95 expression (Fig. 5a) and GluN2A levels (Fig. 5b). GluN2B levels were not affected by either the Aβ_25-35_ or FENM treatment (Fig. 5c). Consequently, the FENM treatment restored the GluN2A/GluN2B ratio (Fig. 5d) GluN2D levels were also analyzed but no effect of the Aβ_25-35_ peptide nor FENM were measured in the synaptosomal preparations (Fig. 5e). The levels of phosphorylated GluN2A/B subunits were analyzed to get an indirect idea of their respective activations. P(Tyr^1325^)-GluN2A was non-significantly decreased after Aβ_25-35_ injection in hippocampal synaptosomes (–26%, Fig. 5f) and no significant effect of FENM was observed in comparison with Aβ_25-35_ (Fig. 5f). On the contrary, P(Ser^1303^)-GluN2B was significantly increased after Aβ_25-35_ injection (+34%, Fig. 5g) and the FENM treatment prevented the increase (Fig. 5g), suggesting that the drug more efficiently restored alteration in GluN2B subunits as compared to GluN2A.

**Figure 5.**
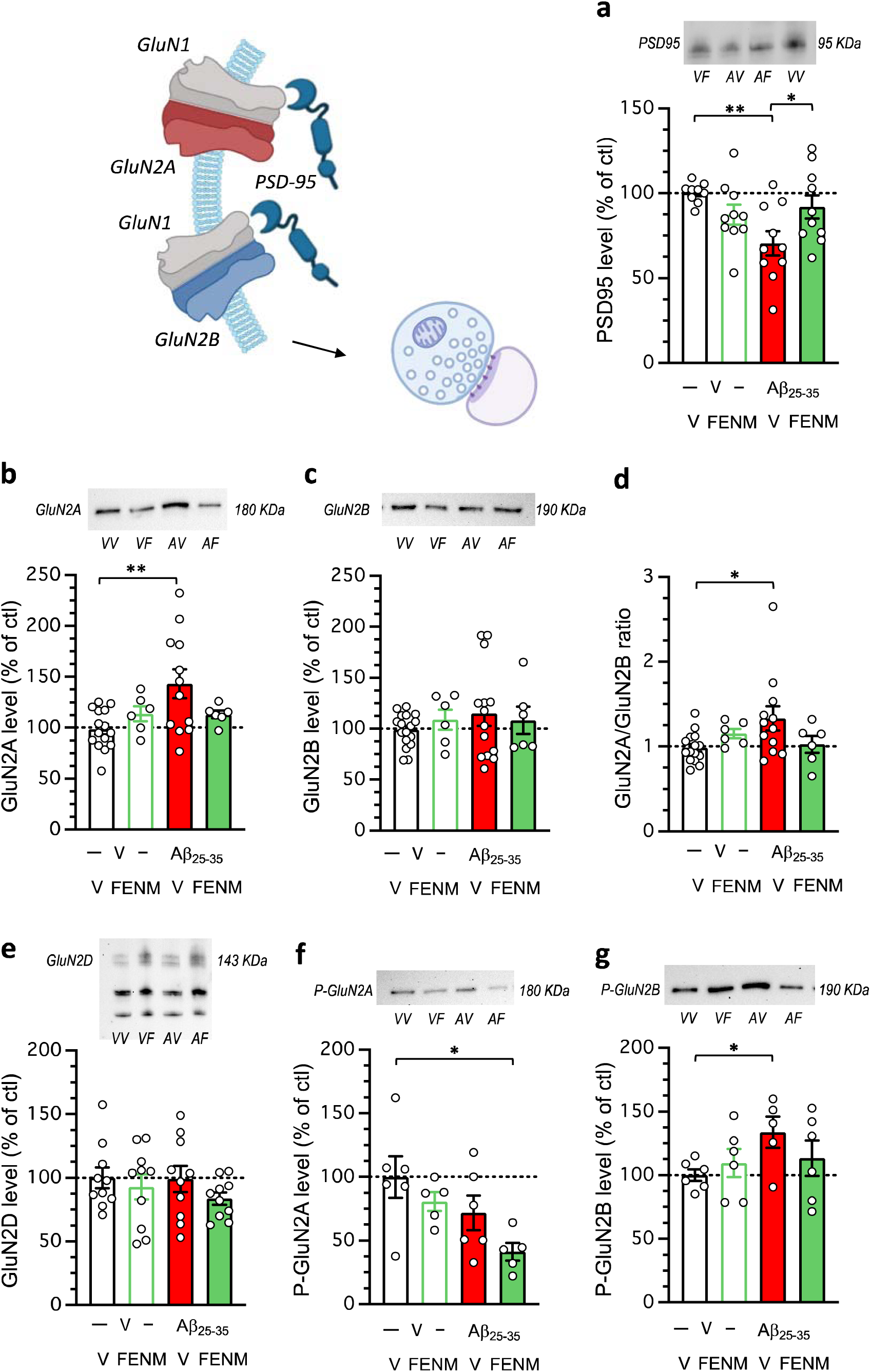
Protective effects of a SC infusion of FENM (0.1 mg/kg/day), on Aβ_25-35_-induced alteration of PSD-95 and NMDAR subunits expressions in synaptosome preparations of mouse hippocampus: (**a**) PSD-95, (**b**) GluN2A, (**c**) GluN2B, (**d**) GluN2A/GluN2B ratio, (**e**) GluN2D, (**f**) P(Tyr^1325^)-GluN2A, (**g**) P(Ser^1303^)-GluN2B. Animals were sacrificed 7 days after the peptide injection. Typical blots are shown above each graph. The whole immunostained and and stain-free blots are shown in Supplementary Figures 3 and 4. The number of animals per group was N = 5-17 per group. * *p* < 0.05, ** *p* < 0.01; Dunnett’s test. Illustrations were done with www.biorender.com.

### FENM treatment by SC infusion or repeated IP injections improved spatial memory impairment in APP/PS1 mice

Results in the Aβ_25-35_ AD model, indicated that the most active doses were 0.3 mg/kg for IP injection, although it must be noted that the object recognition test but not spontaneous alternation, confirmed here the previous results by Couly et al. [31], and 0.1 mg/kg/day for SC infusion. These dosages were therefore selected for drugs administration over 4 weeks period in 10-month-old APP/PS1 mice (Fig. 6a). Animal body weight was monitored daily after osmotic pump implantation or during repeated drug administration, WT and APP/PS1 mice exhibited stable body weight throughout the 4 weeks of treatment (Fig. 6b-d). The repeated IP FENM treatment did not affect the animals’ weight, while the SC infusion treatment had a positive impact for WT and APP/PS1 mice (Fig. 6d-b). Indeed, the average daily weight gain was significantly higher than zero for both FENM-infused WT and APP/PS1 groups, contrarily to controls and FENM IP injected groups (Fig. 6d).

**Figure 6.**
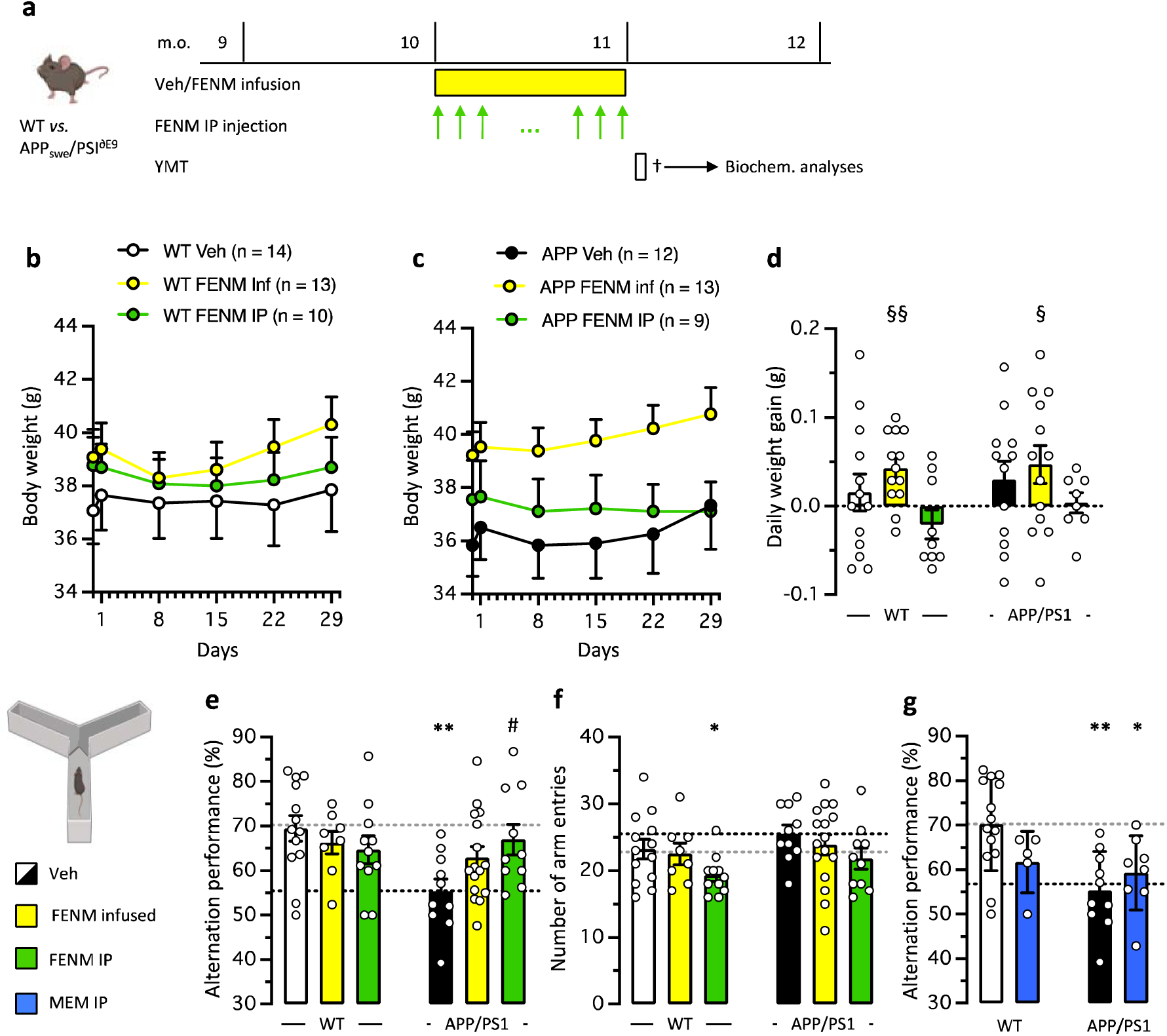
(**a**) Experimental protocol. Wildtype (WT) or APP /PS1^∂E9^ (APP/PS1) mice were infused SC with vehicle solution (Veh) or FENM (0.1 mg/kg/day) or repeatedly injected IP with FENM (0.3 mg/kg) or MEM (0.3 mg/kg) between 10 to 11 month-of-age (m.o.). Surgery was performed on day 1, infusion lasted 28 days, and a Y-maze test (YMT) was performed on day 30. On day 31, animals were sacrificed (†) before biochemical analyses. (**b**,**c**) body weight measured weekly for WT (b) and APP/PS1(c) treated groups. (**d**) Daily weight gain between day 1 to day 29. § p < 0.05, §§ p < 0.01 *vs*. 0 (one-column *t*-test). (**e**) Spontaneous alternation performances and (**f**) total numbers of arm entries for FENM-treated groups and (**g**) spontaneous alternation performances for MEM-treated groups, in the Y-maze test. * *p* < 0.05, ** *p* < 0.01 *vs*. Veh-treated WT group; # p < 0.05 *vs.* Veh-treated APP/PS1 group; Dunnett’s test.

The protective effect of FENM on spatial memory impairment in APP/PS1 mice was first determined through behavioral testing, after the 4 weeks of treatments. Animals were tested for spontaneous alternation in the Y-maze, a rapid measure of hippocampus-dependent spatial working memory in rodents. Veh-treated (control) APP/PS1 mice showed a significant decrease in alternation compared to control WT animals (Fig. 6e). FENM infusion attenuated the deficit (the group performance was not significantly different from control WT value) while repeatedly injected FENM led to a significant prevention of the deficit in APP/PS1 mice (Fig. 6e). Interestingly, the number of arm entries was not different between Veh-treated WT and APP/PS1 groups or following FENM infusion but decreased following repeatedly injected FENM (significantly for WT mice) (Fig. 6f). MEM was also repeatedly injected IP in APP/PS1 mice and WT controls (Fig. 6g) at the active dose of 0.3 mg/kg (Fig. 1i). The drug failed to attenuate the spontaneous alternation impairment in APP/PS1 mice (Fig. 6g).

### FENM treatment by SC infusion or repeated IP injections attenuated neuroinflammation and amyloid load in APP/PS1 mice

Neuroinflammation was evaluated in the hippocampus by measuring the level of TNFα and IL-6. TNFα levels were significantly increased in the hippocampus of control APP/PS1 mice as compared with control WT mice and this increase was attenuated by infused FENM and significantly prevented by repeatedly injected FENM (Fig. 7a). IL-6 levels were significantly increased in control APP/PS1 mice as compared to control WT animals and this increase was significantly prevented by both the FENM infusion or repeated injections (Fig. 7b).

**Figure 7.**
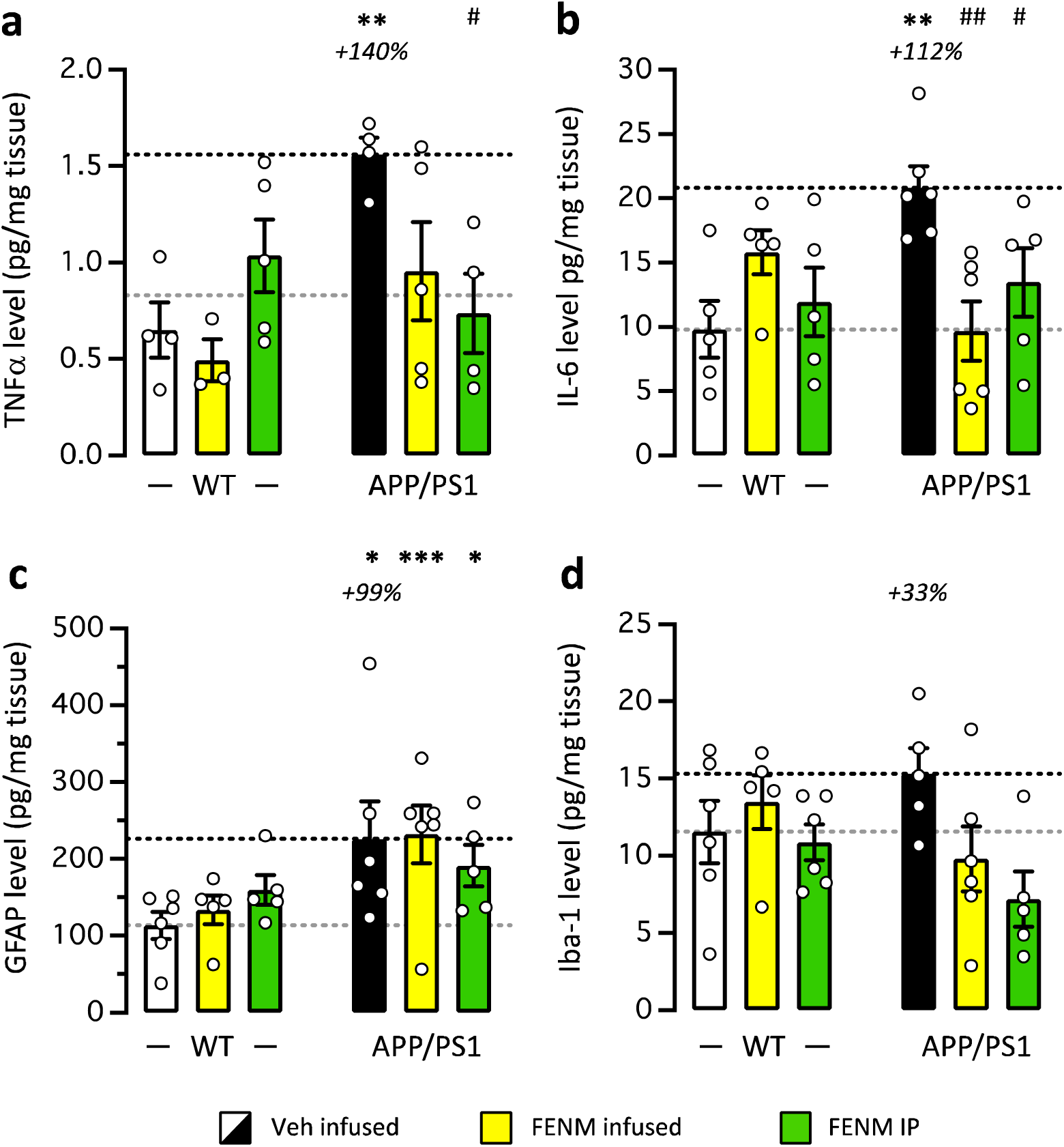
Protective effects of FENM on neuroinflammation in the hippocampus of APP/PS1 mice. Vehicle solution (Veh) or FENM (0.1 mg/kg/day) was infused for 4 weeks SC or FENM (0.3 mg/kg) was injected IP o.d. for 4 weeks. animals were sacrificed 48 h after the last administration day. Levels of the cytokines TNFα (**a**) or IL-6 (**b**) and the glial markers GFAP (**c**) or Iba-1 (**d**) were analyzed in homogenates by ELISA. Data were expressed as percentage of the control, Veh-treated WT group. The number of animal per group was n = 5-6. * *p* < 0.05, ** *p* < 0.01, *** *p* < 0.001 *vs*. the Veh-treated WT group; # *p* < 0.05, ## *p* < 0.01 *vs*. the Veh-treated APP/PS1 group; Dunnett’s test.

GFAP levels measured by Elisa were significantly increased in the hippocampus of control APP/PS1 mice compared to control WT mice (Fig. 7c). Neither the infused FENM nor repeatedly injected FENM decreased GFAP levels (Fig. 7c), suggesting that the treatments marginally affected astroglial reaction after 1 month treatment. IBA-1 level was moderately increased in control APP/PS1 mice as compared to WT (Fig. 7d), but both FENM treatments tended to decrease IBA-1 level in APP/PS1 mice (Fig. 2d).

In the cortex of WT and APP/PS1 mice, we analyzed Bax and Bcl-2 levels. Bax levels were moderately but significantly increased in control APP/PS1 mice and both FENM treatments prevented the increase (Fig. 8a). Bcl-2 levels were unchanged in APP/PS1 mice and whatever the treatment (Fig. 8b).

**Figure 8.**
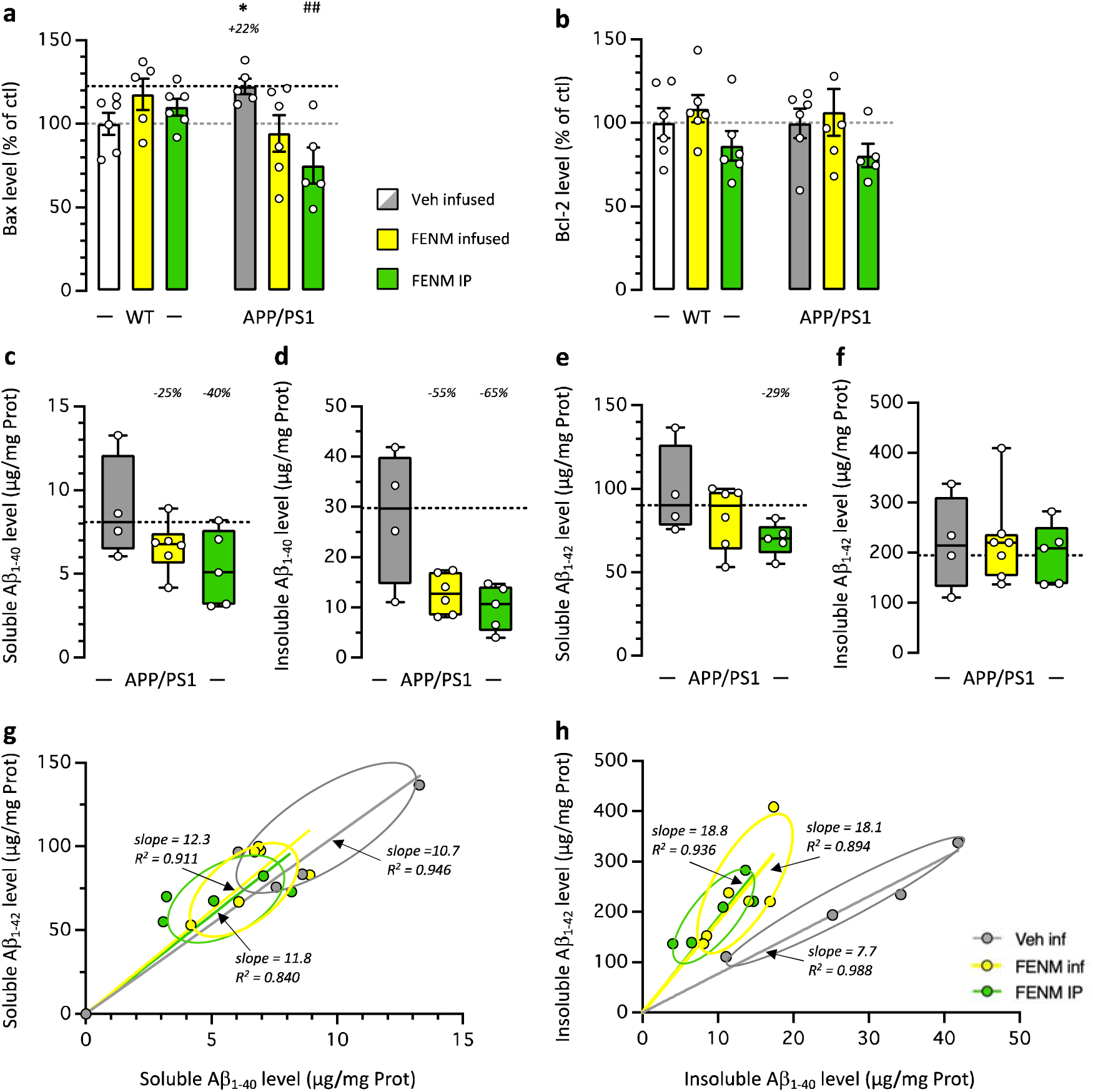
Protective effects of FENM on neuroinflammation in the cortex of APP/PS1 mice. Vehicle solution (Veh) or FENM (0.1 mg/kg/day) was infused for 4 weeks SC or FENM (0.3 mg/kg) was injected IP o.d. for 4 weeks. animals were sacrificed 48 h after the last administration day. Levels of the pro-apoptotic marker Bax (**a**), the anti-apoptotic marker Bcl-2 (**b**), and the soluble (**c**, **e**) and insoluble (**d**, **f**) contents in Aβ_1-40_ (**c**, **d**) and Aβ_1-42_ (**e**, **f**) were analyzed in homogenates by ELISA. Correlations between Aβ_1-40_ and Aβ_1-42_ levels are shown in (**g**) for soluble extracts and in (**h**) for insoluble extracts. Data in (a, b) were expressed as percentage of the control, Veh-treated WT group. Data in (c-f) were expressed as pg/mg of tissue and represented as box-and-whiskers showing the median and range. The number of animal per group was n = 4-6. * *p* < 0.05 *vs*. the Veh-treated WT group; ## *p* < 0.01 *vs*. the Veh-treated APP/PS1 group; Dunnett’s test.

Amyloid load was analyzed in the mouse cortex by quantifying both guanidine-soluble and insoluble forms of Aβ_1-40_ and Aβ_1-42_. Contents in Aβ_1-40_ and Aβ_1-42_ proteins were negligeable in the cortex of WT mice (< 0.15 µg/mg Prot. measured for both Aβ species). However, significant amounts were measured in the cortex of APP/PS1 mice (Fig. 8c-f). The FENM treatments tended to decrease soluble Aβ_1-40_ contents (Fig. 8c) and insoluble Aβ_1-40_ levels (*p* = 0.114 for infused FENM and *p* = 0.064 for repeatedly injected FENM, Mann-Whitney’s test; Fig. 8d), and soluble Aβ_1-42_ contents (*p* < 0.05 for repeatedly injected FENM; Fig. 8e) but did not change insoluble Aβ_1-42_ levels (Fig. 8f). Individual correlations between the levels in Aβ_1-40_ and Aβ_1-42_ confirmed that the FENM treatments decreased similarly both species in soluble forms, with no change in the representation slope (Fig. 8g) while for insoluble forms, the decreases were accompanied with a change in the slopes confirming that the treatments differentially altered Aβ species in all animals (Fig. 8h).

### FENM treatment by SC infusion or repeated IP injections ameliorated LTP impairment in the hippocampal DG in APP/PS1 mice

Based on the FENM effects on NMDAR subunits seen in Aβ_25-35_ mice and to investigate the synaptic mechanism underlying impairments in spatial learning and memory, we examined the hippocampal long-term potentiation (LTP) in APP/PS1 mice. The field excitatory postsynaptic potential (fEPSP) of the CA1 was evoked by stimulating the Schaffer collateral in CA2 in acute hippocampal slices (Fig. 9a). For LTP induction, we applied a high frequency stimulation (HFS, 100 Hz for 1 s), a well-established protocol known for robustly inducing LTP [51]. HFS was delivered to brain slices from both WT and APP/PS1 mice following a 10-min stable baseline at 15 s intervals. Typical fEPSP traces from different groups are shown in Fig. 9b. The fEPSP amplitude was normalized to the average fEPSP amplitude during the baseline (Fig. 9c-e). The average fEPSP amplitude during the last 10 min after HFS (70–80 min) were compared to assess LTP induction and maintenance, respectively (Fig. 9f). In WT mice, the amplitude of fEPSP increased immediately after HFS and then regularly during the measurement period, indicating successful LTP induction and maintenance (Fig. 9c). The average fEPSP amplitude was also increased after HFS in APP/PS1 mice (the first 2 min: 139.4 ± 8.9% for WT mice *vs.* 124.4 ± 7.3% for APP/PS1 mice), but gradually decreased with time (the first 2 min: 137.0 ± 5.9% for WT mice *vs.* 114.0 ± 0.8% for APP/PS1 mice), showing impaired LTP maintenance in APP/PS1 mice. Indeed, a significant decrease in normalized slope fEPSP of APP/PS1 was measured in comparison of WT during the last 10 min (Fig. 9f). Both FENM treatments did not affect maintenance of fEPSP amplitudes in WT mice (Fig. 9d) but allowed a recovery of fEPSP amplitudes in APP/PS1 mice (Fig. 9e), indicating that the drug prevented fEPSP impairment in APP/PS1 mice.

**Figure 9.**
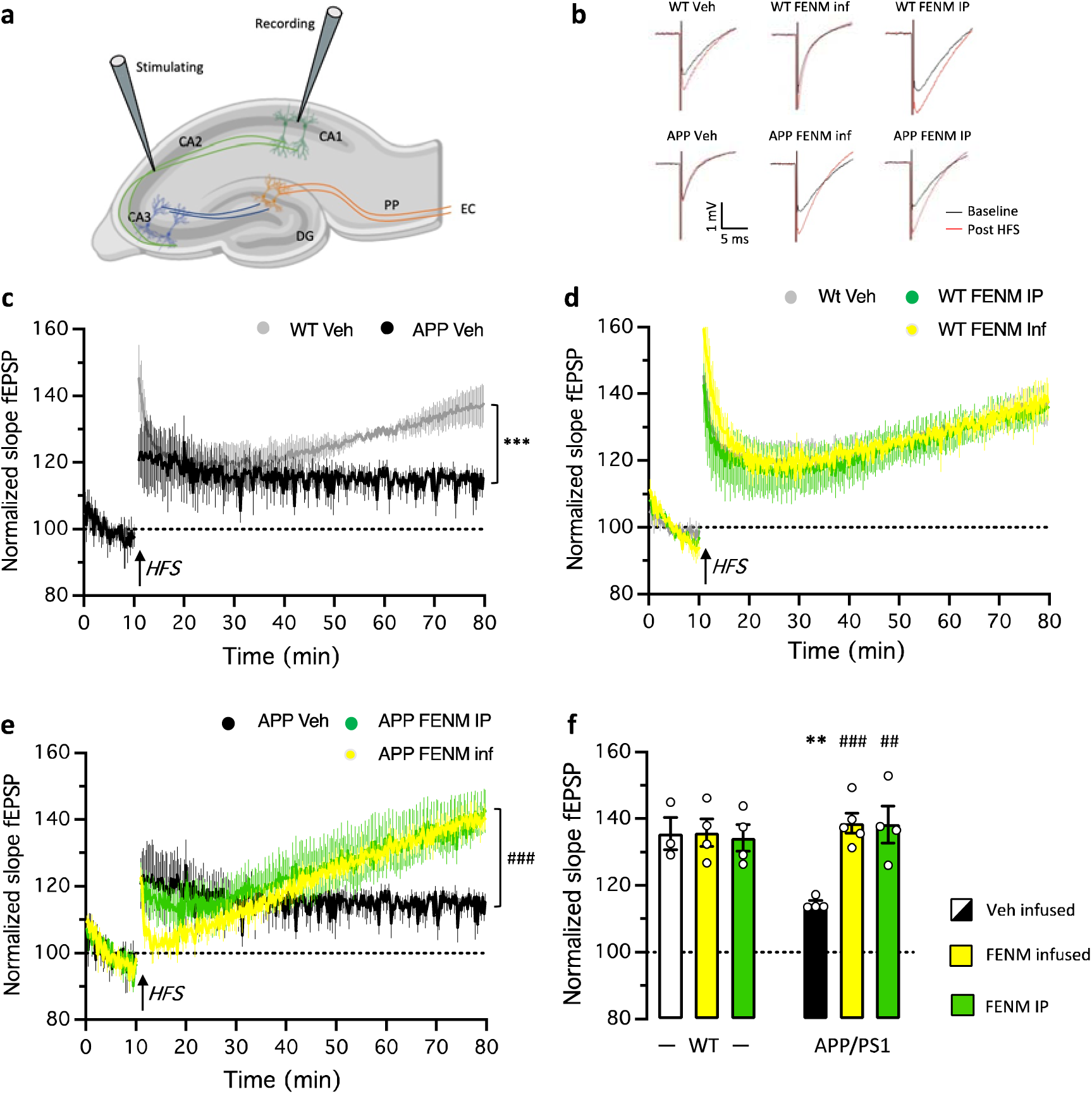
Recovery of LTP in the hippocampal dentate gyrus in APP/PS1 mice by FENM treatments. (**a**) Schematic illustration of field excitatory postsynaptic potential (fEPSP) recordings in hippocampal slices. Abbreviations: CA1∼3, *Cornus Ammona* 1∼3 layers; DG, dentate gyrus; EC, entorhinal cortex; PP, perforant path. (**b**) Typiical examples of the fEPSPs recorded in the DG region in acute mouse hippocampal slices before (black) and 60 min after (red) high frequency stimulations (HFS). (**c**-**e**) The mean amplitude of fEPSPs, expressed as a percentage of the baseline level, is plotted for: (c) the Veh-treated WT and APP/PS1 groups; (d) the Veh-treated, FENM-infused and FENM repeatedly IP injected WT groups; and (e) the Veh-treated, FENM-infused and FENM repeatedly IP injected APP/PS1 groups. It shows the 10 min of baseline recording and 70 min of post⍰HFS (arrow) recording. Responses were evoked and collected every 15 s. (**f**) Average amplitude of fEPSPs during the the last 10 min post⍰HFS recorded in WT and APP/PS1 mice. Data are presented as the mean ± SEM, n = 4-5 animals per group, ** *p* < 0.01, *** *p* < 0.001 *vs*. Veh-treated WT group; ## *p* < 0.01, ### *p* < 0.001 *vs*. Veh-treated APP/PS1 group; repeated-value one-way ANOVA in (c, e), Mann-Whitney test in (f).

## Discussion

In the present study, we confirmed the neuroprotective efficacy of FENM and demonstrated the efficacy of SC infusion in two mouse models of AD, namely the pharmacological Aβ_25-35_ model and the transgenic APP/PS1 line.

In the first model, FENM was previously reported to prevent Aβ_25-35_ mouse-induced toxicity after repeated daily IP injection [32], with a better and more robust prevention of neuroinflammation as compared with MEM, besides a similar efficacy on cognitive deficits and oxidative stress markers [32]. We here confirmed that infused FENM resulted, at all dosages tested from 0.03 to 0.3 mg/kg/day, in preservation of spatial working memory (spontaneous alternation) and recognition memory (novel object test), two memory processes generally altered in AD models. The dose-response profiles appeared at a similar dosage but exhibited more sustained protection than the repeated IP administration of either FENM or MEM in both tests. The most active dosage, 0.1 mg/kg/day, was then used for morphological and biochemical assays. FENM infusion confirmed the ability of the drug to strongly prevent inflammation in the hippocampus, both in terms of astroglial response (GFAP immunolabelling), microglial reaction (IBA-1 immunolabelling), and cytokines release after Aβ_25-35_ insult. Notably, infusion of FENM at 0.1 mg/kg/day was the most active dose to prevent TNFα and IL-6 increase. Both cytokines are mainly released by reactive astroglial and microglial cells. Our biochemical analyses are thus consistent with the effects we measured for FENM on glial cell reactivity.

The impact of FENM infusion on oxidative stress was also analyzed in terms of the levels of peroxidation of the membrane lipids and nitrosylation of proteins in the cortex and hippocampus, respectively. All doses tested significantly prevented the increases in lipid peroxidation and protein nitrosylation. NMDARs have a dual effect on cellular redox level. First, Na^+^ and Ca^2+^ ions entering via over-activated NMDAR, as seen in neurodegeneration, can rapidly overwhelm the entire mitochondrial respiratory capacity, leading to leak of electronic integrity of the respiratory chain and promoting the generation of reactive oxygen species [52]. Second, oxidative stress and NMDAR hypofunction have also been proposed to be related in several physiopathological processes, including schizophrenia [53]. Infusion of FENM was particularly effective in preventing Aβ_25-35_-induced oxidative stress. Indeed, all doses tested prevented lipid peroxidation as well as protein nitrosylation, suggesting that alterations of both mitochondrial respiration or endogenous antioxidant defense systems were alleviated by the NMDAR antagonist.

Another direct consequence of Aβ_25-35_-induced toxicity and synaptic loss we examined was the level of PSD-95 expression. PSD-95 is an accurate marker of postsynaptic degeneration that present altered localization in human AD patient brains [54], and decreased expression in advanced stages of neurodegeneration in amyloid transgenic mouse models [53,54], and in Aβ infused pharmacological models [55,56]. *In vitro*, PSD-95 and Synaptophysin levels are decreased in a concentration– and time-dependent manner after exposure of rat primary hippocampal neurons to Aβ_1-42_ or Aβ_25-35_ [55]. *In vivo*, Aβ_25-35_ *in situ* injection into the hippocampus CA1 layer directly altered PSD-95 and GluN2B in CA1 and the dentate gyrus [56]. It has been proposed that Aβ, by modifying the conformation of the NMDAR C-terminal domain [57] and thus the NMDAR interaction with protein phosphatase 1 (PP1), altered synaptic stability [58]. Here, we confirmed that both in whole hippocampus homogenates and in synaptosomes, *in vivo* administration of Aβ_25-35_ significantly decreased PSD-95 levels. This effect was prevented by the infusion of FENM and can be considered as a direct measure of FENM-induced protection of synapses integrity. The previously demonstrated affinity of FENM on NMDAR GluN subunits at the synapse could thus result in protecting, directly or indirectly, the integrity and the activity of NMDAR against glutamatergic excitotoxicity, avoiding the endocytosis of NMDAR and consequently PSD-95 impairment as observed in AD [59].

NMDAR activity is not only a vector of amyloid toxicity but also a key and subtle actor of the neurodegenerative process. In the AD patient brain, changes in NMDAR subunits expression were described. Using immunohistochemistry and confocal microscopy, Yeung *et al.* [60] recently characterized the expression of NMDAR subunits in *post-mortem* human brain tissue. They described increased level of GluN1 in the *stratum moleculare* and *hilus* of the dentate gyrus (DG), in the *stratum oriens* of the CA2 and CA3, in the *stratum pyramidale* of the CA2, and in the *stratum radiatum* of the CA1, CA2, and CA3 subregions, as well as in the entorhinal cortex. GluN2A expression was significantly increased in AD compared with control in the *stratum oriens*, *stratum pyramidale*, and *stratum radiatum* of CA1 [60], suggesting either an up-regulation of GluN2A expression or a relocalization to the synapse. Indeed, GluN2A NMDARs are mainly pyramidal synaptic receptors, concentrated within the post-synaptic density and scaffolded by PSD-95. They are involved in various form of synaptic plasticity such as long-term potentiation (LTP) [61]. Ca^2+^ entry through GluN2A NMDARs increases cAMP response element binding protein (CREB) and brain-derived neurotrophic factor (BDNF) signaling. An indirect measure of GluN2A activity through the phosphorylation level on Tyr^1325^, showed no difference between Aβ_25-35_ and control mice. However, infused FENM decreased the level of P(Tyr^1325^)-GluN2A suggesting a down regulation of NMDAR activity at the synapse. The drug treatment could thus further decrease GluN2A activity in Aβ_25-35_ mice.

Extrasynaptic NMDAR are predominantly GluN2B-expressing NMDAR and exert different if not opposite actions on post-synaptic neurons [62], in particular by inhibiting CREB signaling [63]. Because the NMDARs are localized closer to the neuronal soma, Ca^2+^ entry resulting from their sustained over-activation due to glutamate spillover from synapses or Aβ-induced alteration of glutamate reuptake by astrocytes, could directly alter mitochondrial membrane potential and result in mitotoxicity and, consequently, cytotoxicity [23]. Further, NMDAR subunits, and particularly the GluN2B subunit, are mobile on the plasma membrane surface between the synapse and extrasynaptic space [64]. These elements led to the consideration that synaptic GluN2A NMDARs are involved in the physiological response to glutamate, including long-term potentiation, and are pro-survival while extrasynaptic GluN2B NMDARs are involved in the neurotoxicity, including long-term depression or spine shrinkage, as observed in excitotoxic events and in neurodegenerative pathologies [65–68]. In Aβ_25-35_ mice, GluN2B levels were unchanged in homogenates and synaptosomes. However, P(Ser^1303^)-GluN2B levels were increased in Aβ_25-35_ mice suggesting that the toxicity increased extrasynaptic NMDAR activity. In the hippocampal neuron cultures of rat, it was shown that the phosphorylation P(Ser^1303^)-GluN2B was increased and revealed NMDAR-induced excitotoxicity [69]. Infusion of FENM restored all changes, suggesting that the drug is effective in alleviating NMDAR dysfunctions both in terms of regulation, *e.g.*, expression and localization, and activity.

It must be noted that NMDAR composition also includes GluN2C, that is poorly expressed in the hippocampus [70], and GluN2D that is present in interneurons and glial cells [71–73]. Although this latter was shown to be unaffected in our study as neither Aβ_25-35_ nor FENM infusion changed GluN2D levels in synaptosomes, we cannot exclude that this subunit plays a particular role in FENM direct action. Indeed, FENM shows a significantly higher selectivity for GluN2C/2D subunits as compared with GluN2A/2B (unpublished data). As Aβ oligomers induces NMDAR dysfunction in astrocytes and alters the phagocytic capacity of microglia [74,75], FENM by a subunit-selective inhibitory action may slow down neuroinflammation and maintain neuronal homeostasis, synaptic transmission, and plasticity. This hypothesis could explain how FENM could impact, indirectly, the GluN2A/B subunit expression and activation in synaptosome through glial cells activity but also how FENM could act on neuroinflammation.

In order to confirm FENM efficacy following a chronic administration treatment in a transgenic AD mouse line, we analyzed FENM effects in APP/PS1 mice. Numerous AD transgenic lines have been used to report the preclinical neuroprotective effect of MEM, including 5XFAD, Tg2576, Tg4-42, 3xTg-AD, APP23, tgCRND8, and APP/PS1 [for a recent review, see 77]. In APP/PS1 mice, IP administration of MEM at 10 mg/kg during 4 weeks improved left/right discrimination in a water version of the T-maze [78]. A 4-month chronic treatment with MEM at 10 mg/kg/day improved object recognition and allowed a decrease in the number of plaques [79]. At 20 mg/kg but by oral gavage during 4 months, the drug improved learning in the Morris water maze, while decreasing APP and Aβ expression [80]. However, at this dose, we previously showed that animal cognition is directly and completely impaired by MEM, diminishing its relevance for pharmacological translation [32]. In the present study, FENM was not only administered at lower doses of 0.1 mg/kg/day or 0.3 mg/kg, which are devoid of direct effect on animal cognition, but also in older and thus more severely affected APP/PS1 mice, 10-11-mo *vs*. 7-8-mo.

At the behavioral level, FENM proved more effective than MEM. FENM successfully restored a correct spatial working memory in mice, whereas MEM showed no effectiveness at the same dosage. We also confirmed that the pathological elevation of TNFα and IL-6 levels in APP/PS1 mice could be alleviated by both FENM treatments. Interestingly, in 10-mo APP/PS1 mice, which exhibit significant amyloid load [43,81] and correlated memory deficits, one-month treatment with FENM notably reduced several markers of amyloid load [82]. More specifically, the drug decreased levels of both guanidine-soluble Aβ_1-40_ and Aβ_1-42_, which correspond to small oligomeric aggregates. These aggregates represent the most toxic amyloid species, causing acute synaptotoxicity and inducing neurodegenerative processes [83,84].

Some studies in transgenic mouse models suggested that Aβ deposits may serve as a sink for soluble Aβ species. Effective therapeutic strategies could thus potentially decrease soluble Aβ oligomer levels and reduce cognitive deficits without impacting or even increasing Aβ plaque levels [85]. Our observation that FENM only marginally affected constituted guanidine-insoluble, *i.e.* fibrillar, Aβ deposits is thus not necessarily a limitation.

Elisa analyses in the hippocampus of 11-mo APP/PS1 detected limited microglial, but strong astrocytic reaction. Interestingly, FENM treatment totally prevented the former while it has only marginal impact on the second. These glial cells express GluN2C or GluN2D NMDARs [72,86]. One hypothesis explaining the observed results could be that FENM affected the overactivation of microglia, or at least regulated microglial activation into prophagocytic or anti-inflammatory phenotypes, by a selective inhibition of GluN2C/2D subunit-expressing NMDARs. Microglia are involved in the immune response, maintenance of homeostasis, extracellular signaling, phagocytosis, antigen presentation and synaptic pruning in the brain and notably, through regulation of neuroinflammation, contribute to AD progression. By regulating microglial reactivity, FENM could improve the clearance of Aβ in brain, as observed in APP/PS1, suggesting a better glutamatergic activity through microglial NMDAR.

The result that GFAP levels was only marginally affected by FENM is surprising as astrocytes play a key role in Glu release and subsequent activation of presynaptic NMDARs, and in Glu reuptake at the synapse. They therefore contribute to the limitation of extrasynaptic NMDARs activation in physiological conditions. Moreover, the astrocytic control of glutamatergic synapse is particularly dependent on the cytokine TNFα that acts as a gating factor for astrocyte activation [87]. The efficacy of FENM treatments to alleviate pathologically increased levels of TNFα without a direct consequence on astroglial activation is therefore paradoxical. Finally, another explanation could be that FENM treatment could have been started late while the astrocyte reaction was well established.

Finally, we observed that both FENM treatments restored LTP in APP/PS1 mice. Synaptic plasticity of the hippocampus is fundamental for hippocampal memory formation, including learning and spatial memory. NMDARs are crucial for synaptic plasticity, LTP induction and maintenance. LTP alteration occurs at 9-15-mo in APP/PS1 mice, *i.e.*, in mature adult transgenic APP/PS1 mice, *vs*. 19-25-mo non-transgenic controls [88]. In the present experiments, APP/PS1 mice still demonstrated some LTP induction, but with a lesser intensity as compared to control animals, and a significant impairment of LTP maintenance over time. Interestingly, the GluN2B subtypes of synaptic NMDARs are key player in the physiological regulation of LTP and resulting plasticity during development [89]. Ji et al. [90] recently described an accelerated shift in NMDAR subtypes from GluN2B to GluN2A to account for the LTP decrease in APP/PS1 animals in relation to the alteration of the hippocampal neurogenesis also observed in these mice [90]. In Aβ_25-35_ mice, we observed that Aβ toxicity increased the GluN2A/GluN2B ratio in hippocampus synaptosomes and that the FENM treatment restored the ratio. The drug had therefore, whatever the treatment protocol, a major impact on hippocampal plasticity, coherent with its ability to improve cognition in APP/PS1 animals.

## Conclusion

The present study demonstrated that FENM is an efficient neuroprotective drug in the Aβ_25-35_ pharmacological mouse model of AD or in APP/PS1 mice, when continuously infused SC. We also described that FENM is neuroprotective after repeated IP injections in APP/PS1 mice while MEM fails to produce effect at doses relevant for pharmacological performance assessment. More specifically, we observed the prevention of memory deficits, neuroinflammation, oxidative stress, apoptosis, alteration in NMDAR subunit expressions in synaptosomes and recovery of LTP maintenance. SC infusion of the drug, an alternative mode of administration that could be implemented using transdermal patch, may offer a progressive and continuous steady-state concentration of the drug, tailored to the needs of AD patient care. Indeed, by developing an appropriate transdermal infusion system, FENM therapy would lead to improved usability, dosage, and compatibility. In particular, the progressive and optimized plasma and brain concentrations of the compound achieved at pharmacologically relevant levels will contribute to the mitigation of adverse side-effect encountered by *per os* repeated dosing.

## Abbreviations

AD: Alzheimer’s disease
APP: amyloid precursor protein
APP/PS1: APP_swe_/PSEN1^∂E9^
Aβ_35_: amyloid-β[25–35] peptide
Bax: Bcl-2–associated X
Bcl-2: B-cell lymphoma 2
DAPI: 4′, 6-diamidino-2-phenylindole
ELISA: enzyme-linked immunosorbent assays
FENM: Fluoroethylnormemantine
fEPSP: postsynaptic field excitation potentials
GFAP: glial fibrillary acidic protein
HFS: high frequency stimulation
HRP: horseradish peroxidase
IBA-1: allograft inflammatory factor 1
ICV: intracerebroventricular
IL-6: interleukin-6
IP: intraperitoneal
LTP: long-term potentiation
MEM: Memantine
mo: month-old
NMDAR: N-methyl-D-aspartate receptor
PBS-T: phosphate buffer saline containing Tween-20
PBS: phosphate buffer saline
PSD-95: postsynaptic density 95 protein
SC: subcutaneous
TBS: Tris-buffered saline
TNFα: tumor necrosis factor-α
Veh: 0.9% saline vehicle solution
WT: wildtype mice

## Acknowledgements

The authors thank the IExplore platform (IGF, Univ Montpellier, CNRS, INSERM, Montpellier, France) for providing part of the WT and APP/PS1 mice used in this study and the CECEMA animal facility (Univ Montpellier, France) for animal handling. Illustrations in figures were prepared using www.biorender.com.

## Authors’ contribution

Conceptualization, TM; Investigation, Data curation, AC, AF, TM; Formal analysis, AC, AF, TM; Resources, AC, AF, HH, TM; Writing, original draft, TM; Writing, review & editing, AC, AF, MV, HH, GR, TM; Funding acquisition, GR, TM; Project administration, TM.

## Availability of data and materials

The datasets and/or analyzed during the current study are available from the corresponding author on reasonable request.

## Ethics approval and consent to participate

Animal procedures were conducted in adherence with the European Union Directive 2010/63 and the ARRIVE guidelines and authorized by the National Ethic Committee (Paris, France): authorization APAFIS #30410-2021031516372048.

## Authors’ details

^1^MMDN, Univ Montpellier, INSERM, EPHE, Montpellier, France. ^2^ReST Therapeutics, Paris, France. ^3^IBMM, Univ Montpellier, CNRS, ENSCM, Montpellier, France. ^4^IGF, Univ Montpellier, CNRS, INSERM, Montpellier, France.

## Supplementary Information

Supplementary material associated with this article can be found, in the electronic version of this article.

---

**Supplementary Table 1.** Commercial kits used in the study.

| *Marker* | *Supplier* | *Reference* | *Batch no.* |
| --- | --- | --- | --- |
| GFAP | Cloud-Clone | SEA068MU | L230517841 |
| IBA-1 | Cloud-Clone | SEC288MU | L230413039 |
| TNFα | Cloud-Clone | SEA133MU | L200923046, L230515648 |
| IL-6 | Cloud-Clone | SEA079MU | L201112013, L230417295 |
| Bax | Cloud-Clone | SEB343MU | L2011131140, L230515329 |
| Bcl-2 | Cloud-Clone | SEA778MU | L201113139, L230517828 |
| Aβ<sub>1-40</sub> | Invitrogen | KHB3481 | 374329-002 |
| Aβ<sub>1-42</sub> | Invitrogen | KHB3441 | 319758-015 |

*Abbreviations:* Bax, bcl-2-like protein 4; Bcl-2, B-cell lymphoma 2, ELISA, enzyme-linked immune-sorbent assay; GFAP, glial fibrillary acidic protein; IBA-1, allograft inflammatory factor 1; IL-6, interleukin-6; TNFα , tumor necrosis factor-α .

**Supplementary Table 2.** Commercial antibodies used in the study.

| *Marker* | *Supplier* | *Description* | *Reference* | *Batch no.* |
| --- | --- | --- | --- | --- |
| *Primary antibodies* |  |  |  |  |
| Iba-1 | Wako | rabbit polyclonal | 019-19741 | PTE0555 |
| GFAP | Sigma-Aldrich | mouse monoclonal | G3893 | 029M4819V |
| GluN2A | Cell Signalling | rabbit polyclonal | #4205 | 3 |
| GluN2B | Cell Signalling | rabbit polyclonal | #4207 | 2 |
| GluN2D | Abcam | rabbit polyclonal | ab35448 | 851239 |
| GluN2A (phospho Y1325) | Abcam | rabbit monoclonal | ab302930 | 1025308-3 |
| GluN2B (phospho S1303) | Abcam | rabbit monoclonal | ab81271 | GR221673-20 |
| PSD95 [6G6-1C9] | Abcam | mouse monoclonal | ab2723 | GR3248435-12 |
| Nitrotyrosine, clone 1A6 | Sigma-Aldrich | mouse monoclonal | 05-233 | 3403470 |
| *Secondary antibodies* |  |  |  |  |
| Cy3 AffiniPure | Jackson ImmunoResearch | donkey anti-mouse IgG (H+L) | 715-165-150 | 140779 |
| Alexa Fluor 488 AffiniPure | Jackson ImmunoResearch | donkey anti-rabbit IgG (H+L) | 711-545-152 | 140350 |
| HRP-linked antibody | Cell Signalling | anti-rabbit IgG | #7074 | 29 |
| HRP-linked antibody | Abcam | goat anti-mouse IgG H&L | ab6789 | GR3395730-9 |

**Abbreviations:** Iba-1, Ionized calcium-binding adapter molecule 1; GFAP, glial acidic fibrillary protein; GluN2A~D, glutamatergic N-methyl-D-aspartate receptor 2A~D subunit; PSD95 [6G6-1C9], Postsynaptic density protein 95; Cy3, Cyanine Cy3.

**Supplementary Table 3.**
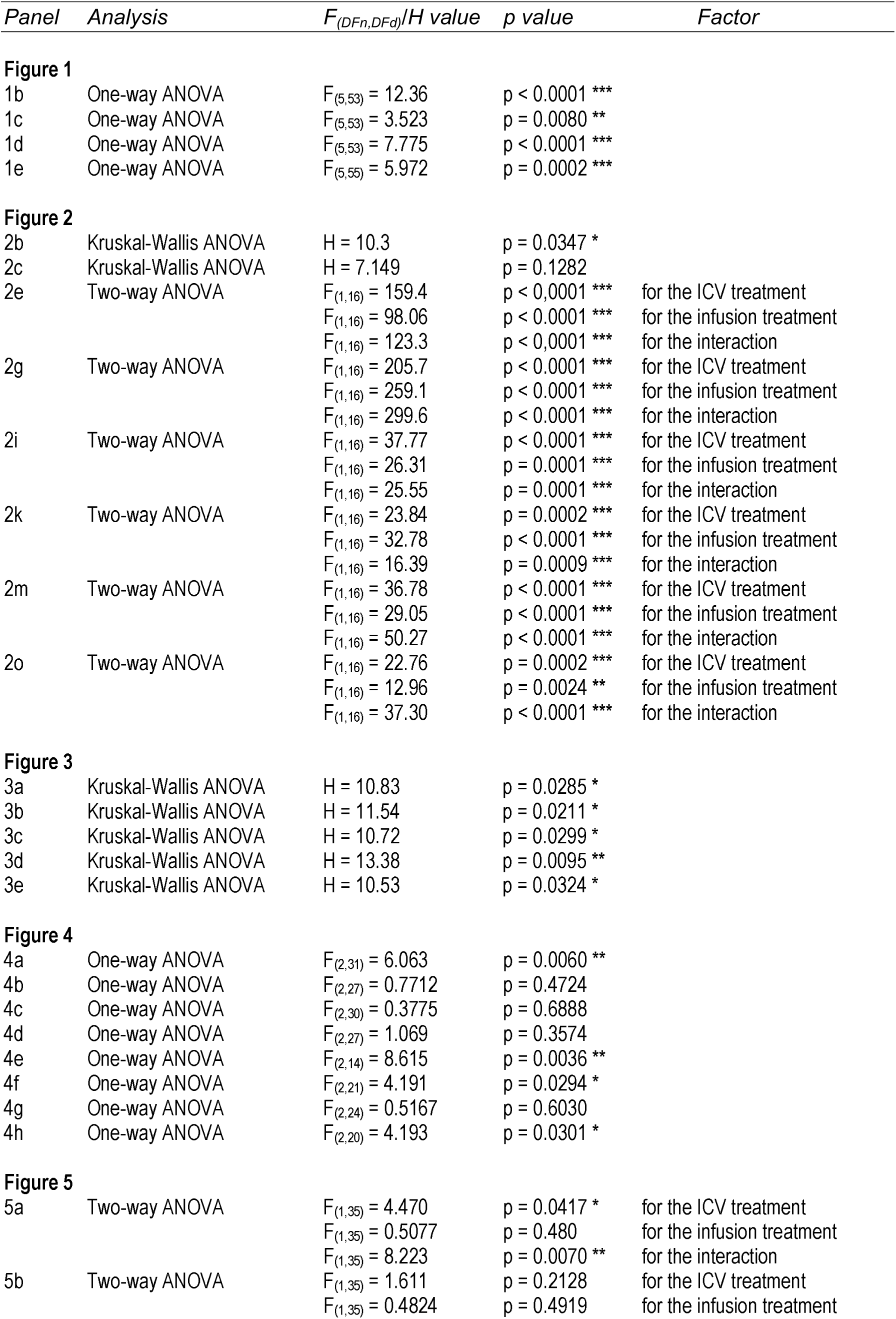
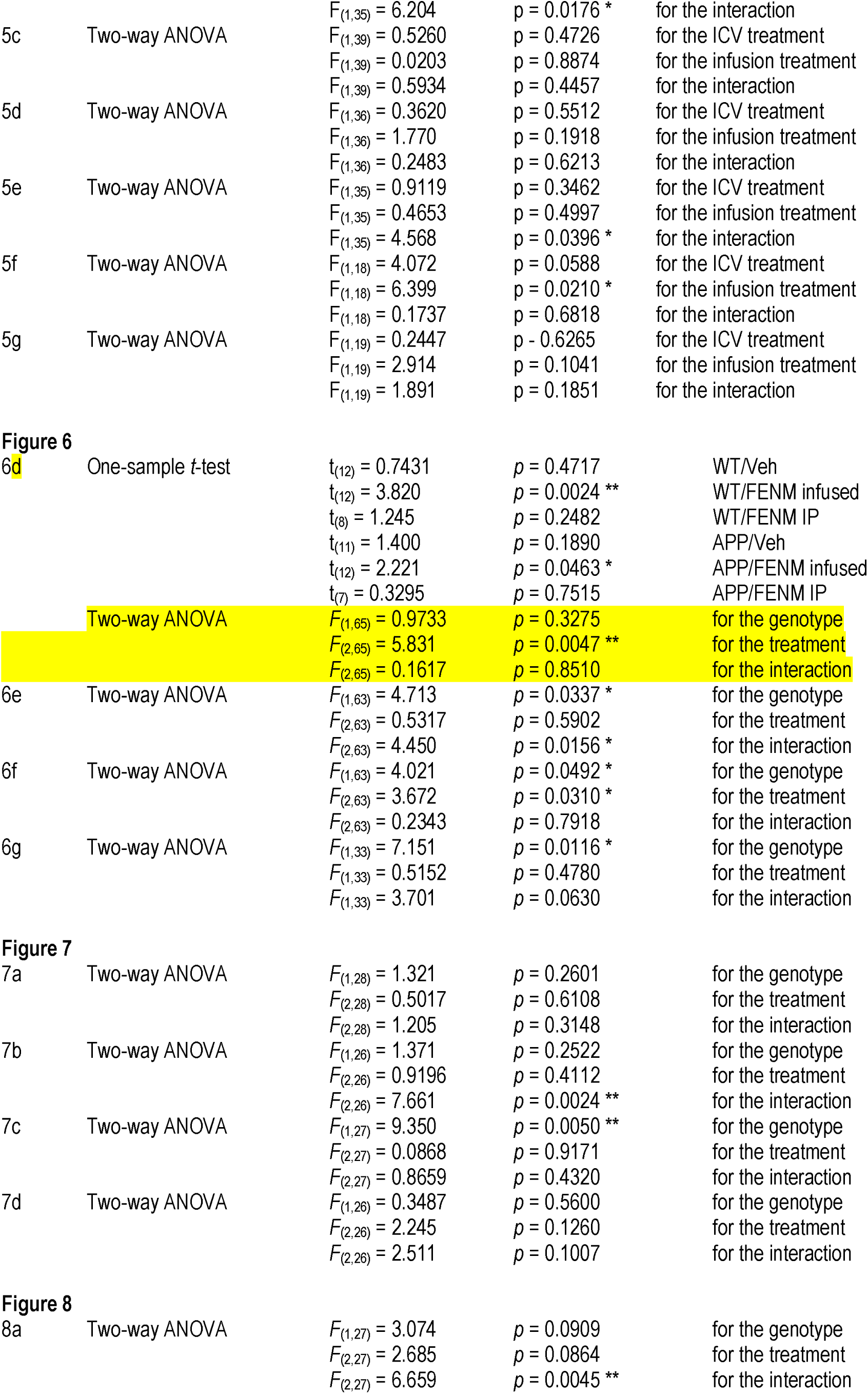
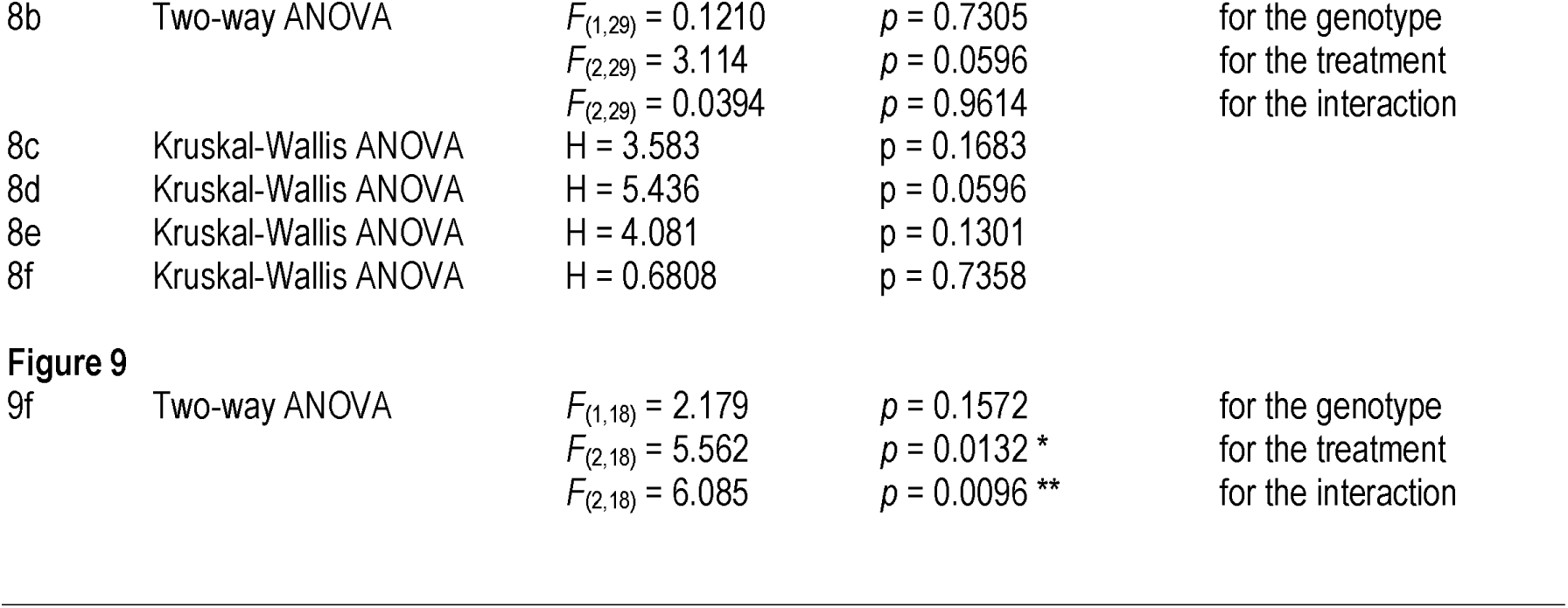
Statistical analyses of variances for the Figures.

**Supplementary Figure 1.**
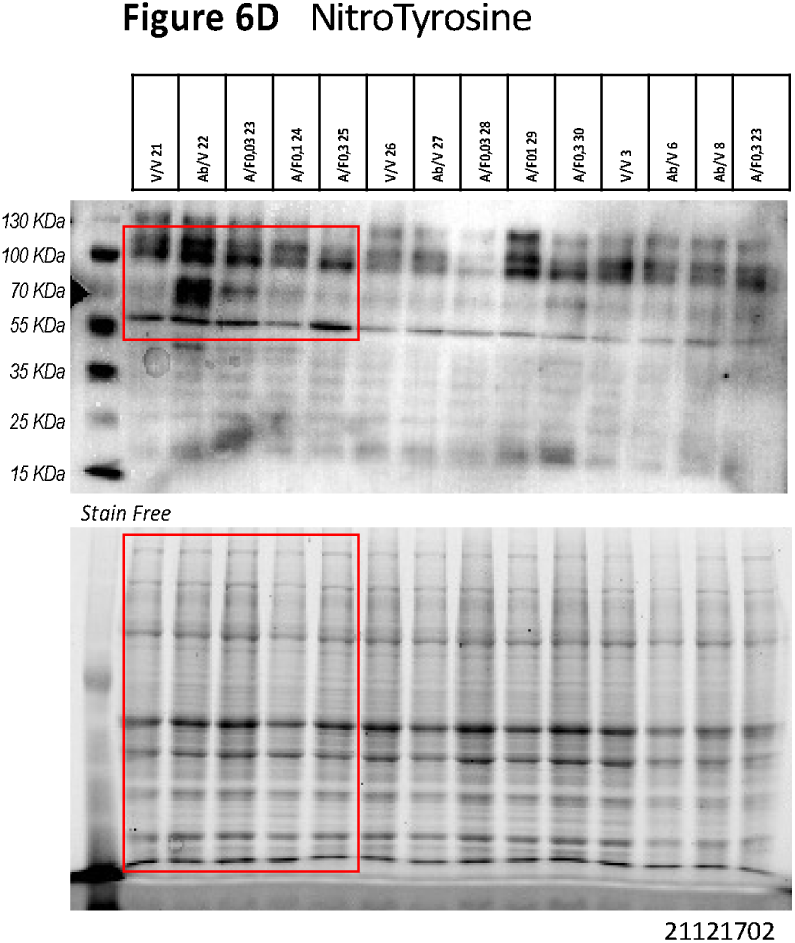

**Supplementary Figure 2.**
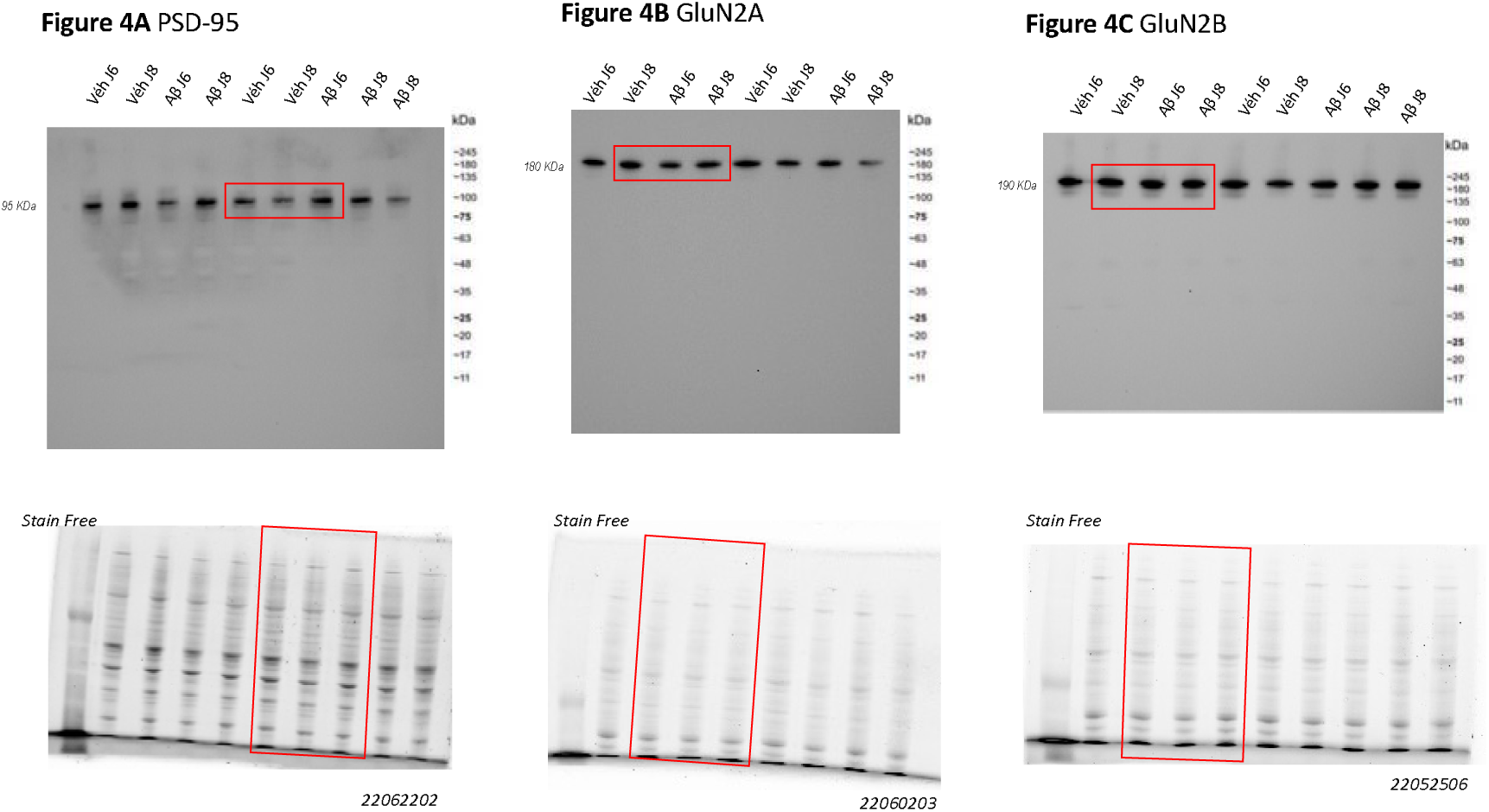

**Supplementary Figure 3.**
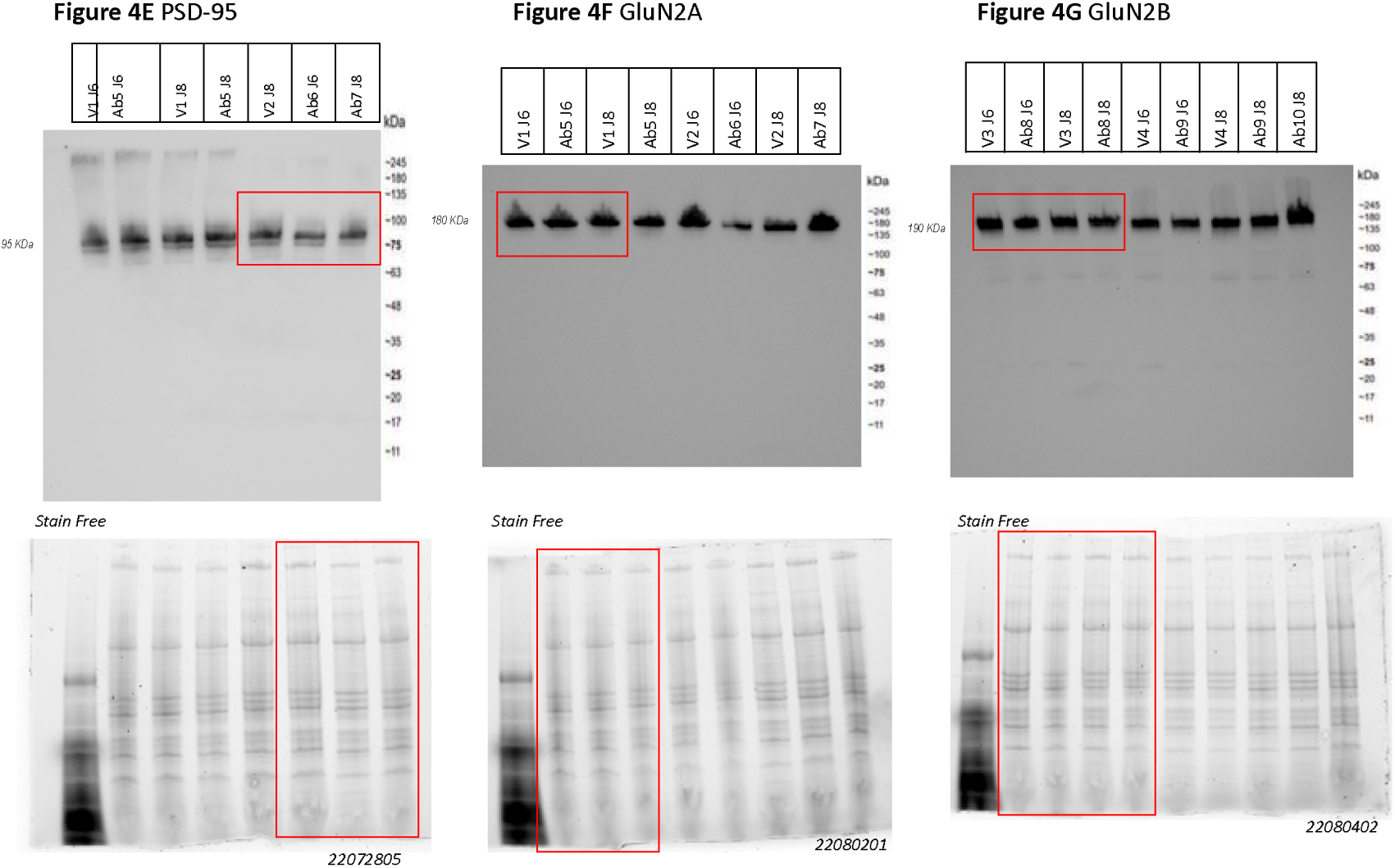

**Supplementary Figure 4.**
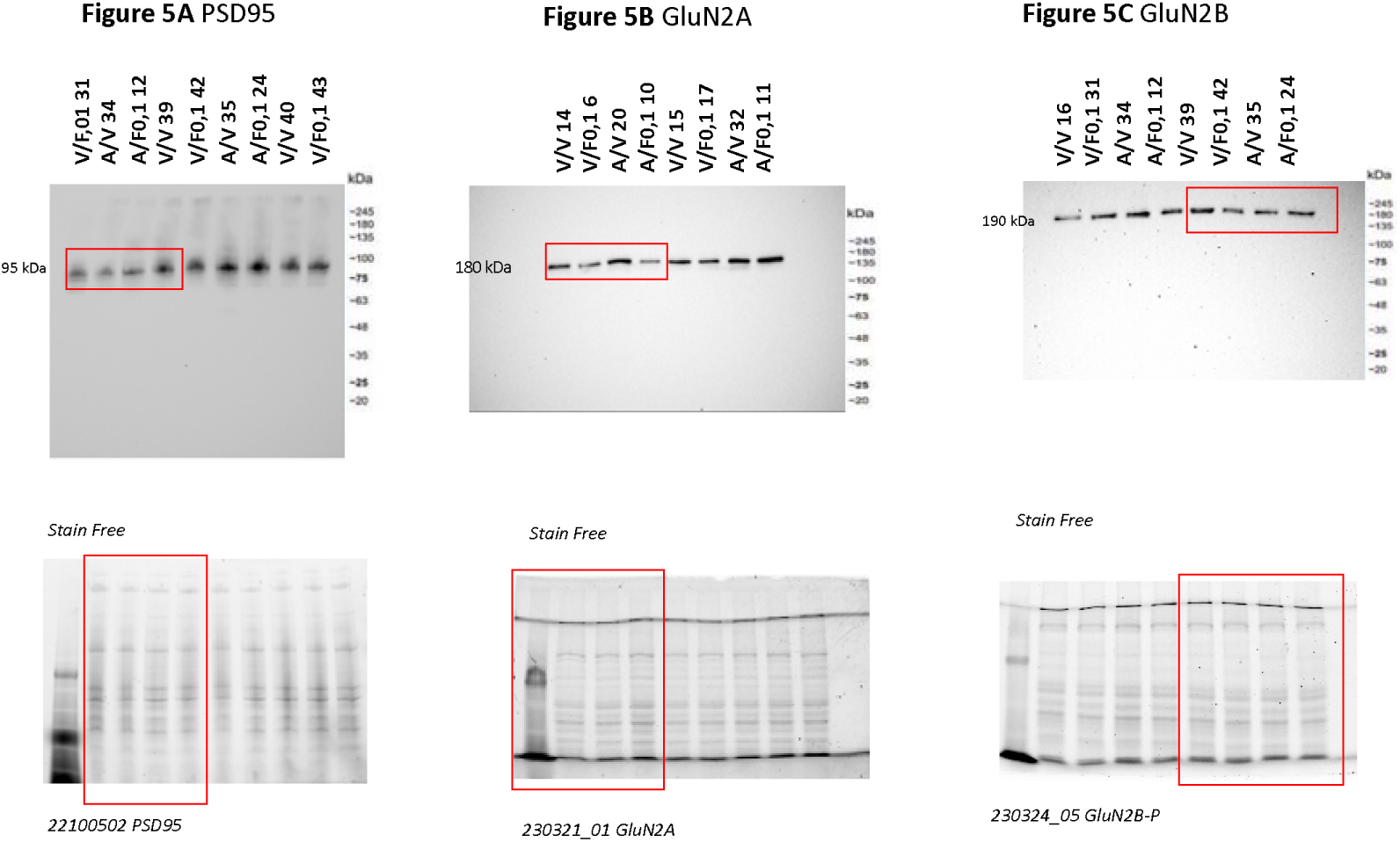

**Supplementary Figure 5.**
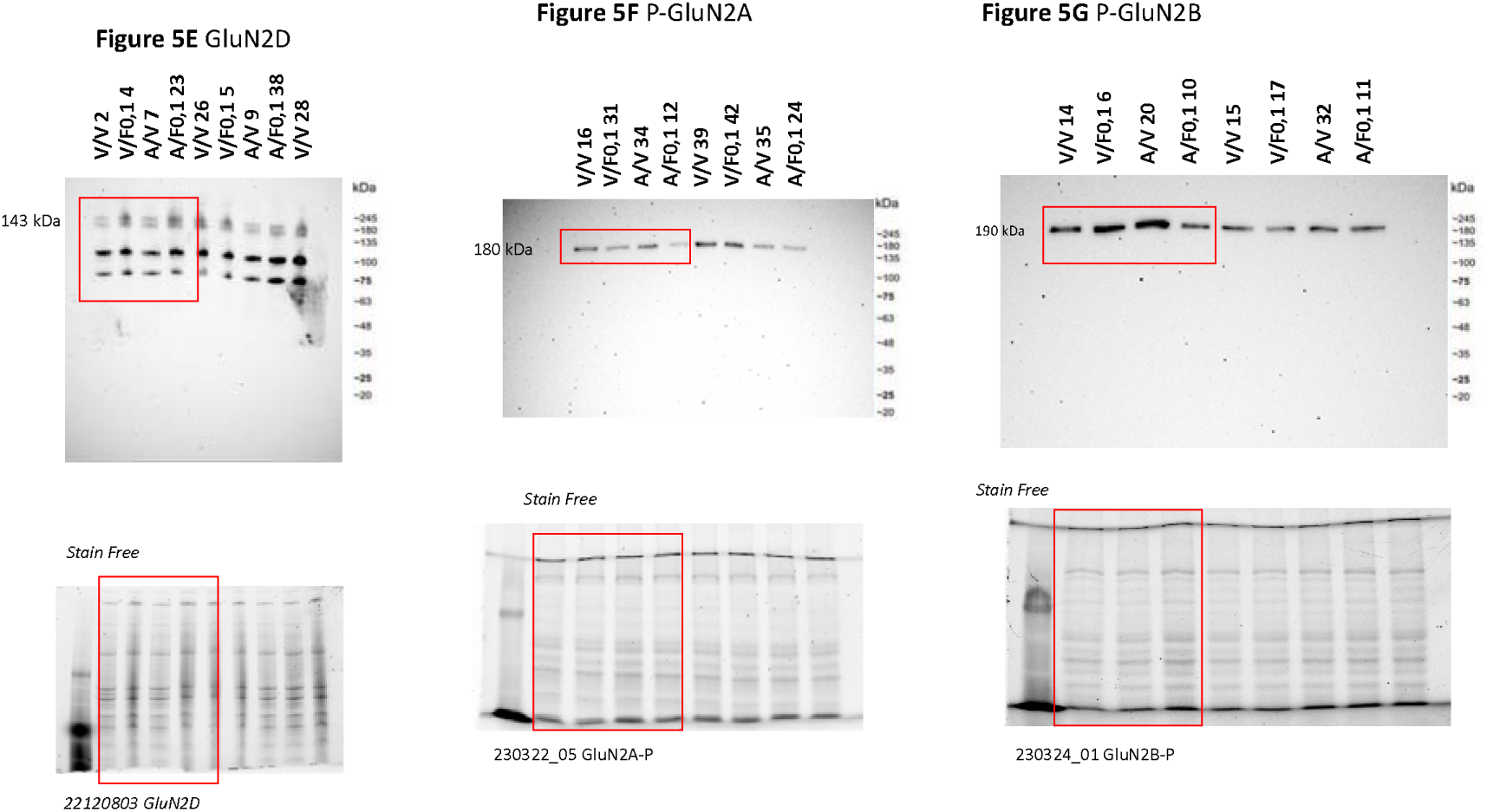

## References

1. Brookmeyer R, Gray S, Kawas C (1998) Projections of Alzheimer’s disease in the United States and the public health impact of delaying disease onset. Am J Public Health 88, 1337–42.

2. Lane CA, Hardy J, Schott JM (2018) Alzheimer’s disease. Eur J Neurol 25, 59–70.

3. Selkoe DJ (1991) The molecular pathology of Alzheimer’s disease. Neuron 6, 487–98.

4. Selkoe DJ (2004) Cell biology of protein misfolding: the examples of Alzheimer’s and Parkinson’s diseases. Nat Cell Biol 6, 1054–61.

5. Chen Y, Yu Y (2023) Tau and neuroinflammation in Alzheimer’s disease: interplay mechanisms and clinical translation. J Neuroinflammation 20, 165.

6. Gouilly D, Rafiq M, Nogueira L, Salabert AS, Payoux P, Péran P, Pariente J (2023) Beyond the amyloid cascade: an update of Alzheimer’s disease pathophysiology. Rev Neurol (Paris) S0035-3787(23)00870-6.

7. Ratan Y, Rajput A, Maleysm S, Pareek A, Jain V, Pareek A, Kaur R, Singh G (2023) An insight into cellular and molecular mechanisms underlying the pathogenesis of neurodegeneration in Alzheimer’s disease. Biomedicines 11, 1398.

8. Frost B, Jacks RL, Diamond MI (2009) Propagation of tau misfolding from the outside to the inside of a cell. J Biol Chem 284, 12845–52.

9. Butterfield DA, Halliwell B (2019) Oxidative stress, dysfunctional glucose metabolism and Alzheimer disease. Nat Rev Neurosci 20, 148–160.

10. Bloom GS (2014) Amyloid-β and tau: the trigger and bullet in Alzheimer disease pathogenesis. JAMA Neurol 71, 505–8.

11. Avila J, Llorens-Martín M, Pallas-Bazarra N, Bolós M, Perea JR, Rodríguez-Matellán A, Hernández F (2017) Cognitive decline in neuronal aging and Alzheimer’s disease: role of NMDA receptors and associated proteins. Front Neurosci 11, 626.

12. Wang R, Reddy PH (2017) Role of glutamate and NMDA receptors in Alzheimer’s disease. J Alzheimers Dis 57, 1041–1048.

13. Cotman CW, Geddes JW, Bridges RJ, Monaghan DT (1989) N-methyl-D-aspartate receptors and Alzheimer’s disease. Neurobiol Aging 10, 603–5.

14. Bliss TV, Collingridge GL (1993) A synaptic model of memory: long-term potentiation in the hippocampus. Nature 361(6407), 31–9.

15. Talantova M, Sanz-Blasco S, Zhang X, Xia P, Akhtar MW, Okamoto S, Dziewczapolski G, Nakamura T, Cao G, Pratt AE, Kang YJ, Tu S, Molokanova E, McKercher SR, Hires SA, Sason H, Stouffer DG, Buczynski MW, Solomon JP, Michael S, Powers ET, Kelly JW, Roberts A, Tong G, Fang-Newmeyer T, Parker J, Holland EA, Zhang D, Nakanishi N, Chen HS, Wolosker H, Wang Y, Parsons LH, Ambasudhan R, Masliah E, Heinemann SF, Piña-Crespo JC, Lipton SA (2013) Aβ induces astrocytic glutamate release, extrasynaptic NMDA receptor activation, and synaptic loss. Proc Natl Acad Sci USA 110, E2518–27.

16. Sattler R, Tymianski M (2001) Molecular mechanisms of glutamate receptor-mediated excitotoxic neuronal cell death. Mol Neurobiol 24, 107–29.

17. Vesce S, Rossi D, Brambilla L, Volterra A (2007) Glutamate release from astrocytes in physiological conditions and in neurodegenerative disorders characterized by neuroinflammation. Int Rev Neurobiol 82, 57–71.

18. Uttara B, Singh AV, Zamboni P, Mahajan RT (2009) Oxidative stress and neurodegenerative diseases: a review of upstream and downstream antioxidant therapeutic options. Curr Neuropharmacol 7, 65–74.

19. Sims JR, Zimmer JA, Evans CD, Lu M, Ardayfio P, Sparks J, Wessels AM, Shcherbinin S, Wang H, Monkul Nery ES, Collins EC, Solomon P, Salloway S, Apostolova LG, Hansson O, Ritchie C, Brooks DA, Mintun M, Skovronsky DM; TRAILBLAZER-ALZ 2 Investigators (2023). Donanemab in early symptomatic Alzheimer disease: The TRAILBLAZER-ALZ 2 randomized clinical trial. J Amer Med Assoc 330, 512–27.

20. van Dyck CH, Swanson CJ, Aisen P, Bateman RJ, Chen C, Gee M, Kanekiyo M, Li D, Reyderman L, Cohen S, Froelich L, Katayama S, Sabbagh M, Vellas B, Watson D, Dhadda S, Irizarry M, Kramer LD, Iwatsubo T (2023) Lecanemab in early Alzheimer’s disease. N Engl J Med 388, 9–21.

21. Cummings J, Aisen P, Lemere C, Atri A, Sabbagh M, Salloway S (2021) Aducanumab produced a clinically meaningful benefit in association with amyloid lowering. Alzheimers Res Ther 13, 98.

19. Danysz W, Parsons CG (2003) The NMDA receptor antagonist memantine as a symptomatological and neuroprotective treatment for Alzheimer’s disease: preclinical evidence. Int J Geriatr Psychiatry 18:S23–32.

20. Chin E, Jaqua E, Safaeipour M, Ladue T (2022) Conventional versus new treatment: comparing the effects of acetylcholinesterase inhibitors and N-methyl-D-aspartate receptor antagonist with Aducanumab. Cureus 14, e31065.

21. Hardingham GE, Bading H (2011) Synaptic versus extrasynaptic NMDA receptor signalling: implications for neurodegenerative disorders. Nat Rev Neurosci 11, 682–96.

22. Xia P, Chen HS, Zhang D, Lipton SA (2010) Memantine preferentially blocks extrasynaptic over synaptic NMDA receptor currents in hippocampal autapses. J Neurosci 30, 11246–50.

23. Parsons MP, Raymond LA (2014) Extrasynaptic NMDA receptor involvement in central nervous system disorders. Neuron 82 279–93.

24. Ametamey SM, Bruehlmeier M, Kneifel S, Kokic M, Honer M, Arigoni M, Buck A, Burger C, Samnick S, Quack G, Schubiger PA (2002) PET studies of ^18^F-Memantine in healthy volunteers. Nucl Med Biol 29, 227–31.

25. Samnick S, Ametamey S, Leenders KL, Vontobel P, Quack G, Parsons CG, Neu H, Schubiger PA (1998) Electrophysiological study, biodistribution in mice, and preliminary PET evaluation in a rhesus monkey of 1-amino-3-[^18^F]fluoromethyl-5-methyladamantane (^18^F-MEM): a potential radioligand for mapping the NMDA-receptor complex. Nucl Med Biol 25, 323–30.

26. Salabert AS, Fonta C, Fontan C, Adel D, Alonso M, Pestourie C, Belhadj-Tahar H, Tafani M, Payoux P (2015) Radiolabeling of [^18^F]-fluoroethylnormemantine and initial in vivo evaluation of this innovative PET tracer for imaging the PCP sites of NMDA receptors. Nucl Med Biol 42, 643–53.

27. Salabert AS, Mora-Ramirez E, Beaurain M, Alonso M, Fontan C, Tahar HB, Boizeau ML, Tafani M, Bardiès M, Payoux P (2018) Evaluation of [^18^F]FNM biodistribution and dosimetry based on whole-body PET imaging of rats. Nucl Med Biol 59:1–8.

28. Beaurain M, Talmont F, Pierre D, Péran P, Boucher S, Hitzel A, Rols MP, Cuvillier O, Payoux P, Salabert AS (2023) Pharmacological characterization of [^18^F]-FNM and evaluation of NMDA receptors activation in a rat brain injury model. Mol Imaging Biol 25: 692–703.

32. 29. Chen BK, Le Pen G, Eckmier A, Rubinstenn G, Jay TM, Denny CA (2021) Fluoroethylnormemantine, a novel derivative of memantine, facilitates extinction learning without sensorimotor deficits. Int J Neuropsychopharmacol 24, 519–531.

30. Chen BK, Luna VM, Shannon ME, Hunsberger HC, Mastrodonato A, Stackmann M, McGowan JC, Rubinstenn G, Denny CA (2021) Fluoroethylnormemantine, a novel NMDA receptor antagonist, for the prevention and treatment of stress-induced maladaptive behavior. Biol Psychiatr 90, 458–472.

31. Couly S, Denus M, Bouchet M, Rubinstenn G, Maurice T (2021) Anti-amnesic and neuroprotective effects of Fluoroethylnormemantine in a pharmacological mouse model of Alzheimer’s disease. Int J Neuropsychopharmacol 24, 142–157.

32. Maurice T, Lockhart BP, Privat A (1996) Amnesia induced in mice by centrally administered β-amyloid peptides involves cholinergic dysfunction. Brain Res 706, 181– 93.

33. Meunier J, Ieni J, Maurice T (2006) The anti-amnesic and neuroprotective effects of donepezil against amyloid β_25-35_ peptide-induced toxicity in mice involve an interaction with the sigma_1_ receptor. Br J Pharmacol 149, 998–1012.

34. Villard V, Espallergues J, Keller E, Vamvakides A, Maurice T (2011) Anti-amnesic and neuroprotective potentials of the mixed muscarinic receptor/sigma_1_ (σ_1_) ligand ANAVEX2-73, a novel aminotetrahydrofuran derivative. J Psychopharmacol 25, 1101– 17.

35. Rodríguez Cruz Y, Strehaiano M, Rodríguez Obaya T, García Rodríguez JC, Maurice T (2017) An intranasal formulation of erythropoietin (Neuro-EPO) prevents memory deficits and amyloid toxicity in the APPSwe transgenic mouse model of Alzheimer’s disease. J Alz Dis 55, 231–248.

36. Gilda JE, Gomes AV (2013) Stain-Free total protein staining is a superior loading control to β-actin for Western blots. Anal Biochem 440, 186–8.

37. Gürtler A, Kunz N, Gomolka M, Hornhardt S, Friedl AA, McDonald K, Kohn JE, Posch A (2013) Stain-Free technology as a normalization tool in Western blot analysis. Anal Biochem 433, 105–11.

36. Maurice T, Volle JN, Strehaiano M, Crouzier L, Pereira C, Kaloyanov N, Virieux D, Pirat JL (2019) Neuroprotection in non-transgenic and transgenic mouse models of Alzheimer’s disease by positive modulation of σ_1_ receptors. Pharmacol Res 144, 315– 30.

37. Ditzler K (1991) Efficacy and tolerability of memantine in patients with dementia syndrome. A double-blind, placebo controlled trial. Arzneimittelforschung 41, 773–80.

38. Nakamura ZM, Deal AM, Park EM, Stanton KE, Lopez YE, Quillen LJ, O’Hare Kelly E, Heiling HM, Nyrop KA, Ray EM, Dees EC, Reeder-Hayes KE, Jolly TA, Carey LA, Abdou Y, Olajide OA, Rauch JK, Joseph R, Copeland A, McNamara MA, Ahles TA, Muss HB (2023) A phase II single-arm trial of memantine for prevention of cognitive decline during chemotherapy in patients with early breast cancer: Feasibility, tolerability, acceptability, and preliminary effects. Cancer Med 12, 8172–83. doi: 10.1002/cam4.5619

39. Ballard C, Thomas A, Gerry S, Yu LM, Aarsland D, Merritt C, Corbett A, Davison C, Sharma N, Khan Z, Creese B, Loughlin P, Bannister C, Burns A, Win SN, Walker Z; MAIN-AD investigators (2015) A double-blind randomized placebo-controlled withdrawal trial comparing memantine and antipsychotics for the long-term treatment of function and neuropsychiatric symptoms in people with Alzheimer’s disease (MAIN-AD). J Am Med Dir Assoc 16, 316–22. doi: 10.1016/j.jamda.2014.11.002

40. Small G, Dubois B (2007) A review of compliance to treatment in Alzheimer’s disease: potential benefits of a transdermal patch. Curr Med Res Opin 23, 2705–13.

41. Winblad B, Cummings J, Andreasen N, Grossberg G, Onofrj M, Sadowsky C, Zechner S, Nagel J, Lane R (2007) A six-month double-blind, randomized, placebo-controlled study of a transdermal patch in Alzheimer’s disease-rivastigmine patch versus capsule. Int J Geriatr Psychiatry 22, 456–67.

42. Beconi MG, Howland D, Park L, Lyons K, Giuliano J, Dominguez C, Munoz-Sanjuan I, Pacifici R. Pharmacokinetics of memantine in rats and mice. PLoS Curr. 2011 Dec 15;3:RRN1291. doi: 10.1371/currents.RRN1291

43. Jankowsky JL, Fadale DJ, Anderson J, Xu GM, Gonzales V, Jenkins NA, Copeland NG, Lee MK, Younkin LH, Wagner SL, Younkin SG, Borchelt DR (2004) Mutant presenilins specifically elevate the levels of the 42 residue beta-amyloid peptide in vivo: evidence for augmentation of a 42-specific gamma secretase. Hum Mol Genet 13, 159–70.

44. Lalonde R, Dumont M, Staufenbiel M, Strazielle C (2005) Neurobehavioral characterization of APP23 transgenic mice with the SHIRPA primary screen. Behav Brain Res 157, 91–8.

45. Garcia-Alloza M, Robbins EM, Zhang-Nunes SX, Purcell SM, Betensky RA, Raju S, Prada C, Greenberg SM, Bacskai BJ, Frosch MP (2006) Characterization of amyloid deposition in the APP_swe_/PS1^dE9^ mouse model of Alzheimer disease. Neurobiol Dis 24, 516–24.

46. Volianskis A, Køstner R, Mølgaard M, Hass S, Jensen MS (2010) Episodic memory deficits are not related to altered glutamatergic synaptic transmission and plasticity in the CA1 hippocampus of the APP_swe_/PS1^δE9^-deleted transgenic mice model of ß-amyloidosis. Neurobiol Aging 31, 1173–87.

47. Kilkenny C, Browne W, Cuthill IC, Emerson M, Altman DG (2010) NC3Rs Reporting Guidelines Working Group. Animal research: reporting in vivo experiments: the ARRIVE guidelines. Br J Pharmacol 160, 1577–9.

48. Paxinos G, Franklin KB (2019) Paxinos and Franklin’s the mouse brain in stereotaxic coordinates. Academic press.

49. Crouzin N, Baranger K, Cavalier M, Marchalant Y, Cohen-Solal C, Roman FS, Khrestchatisky M, Rivera S, Féron F, Vignes M (2013) Area-specific alterations of synaptic plasticity in the 5XFAD mouse model of Alzheimer’s disease: dissociation between somatosensory cortex and hippocampus. PLoS One 8, e74667.

50. Nicholls DG (2008) Oxidative stress and energy crises in neuronal dysfunction. Ann N Y Acad Sci 1147, 53–60.

51. Lin CH, Lane HY (2019) Early Identification and intervention of schizophrenia: insight from hypotheses of glutamate dysfunction and oxidative stress. Front Psychiatry 10, 93.

52. Chiang TI, Yu YH, Lin CH, Lane HY (2021) Novel biomarkers of Alzheimer’s disease: based upon N-methyl-D-aspartate receptor hypoactivation and oxidative stress. Clin Psychopharmacol Neurosci 19, 423–433.

53. Shao CY, Mirra SS, Sait HB, Sacktor TC, Sigurdsson EM (2011) Postsynaptic degeneration as revealed by PSD-95 reduction occurs after advanced Aβ and tau pathology in transgenic mouse models of Alzheimer’s disease. Acta Neuropathol 122, 285–92.

54. Liu J, Chang L, Roselli F, Almeida OF, Gao X, Wang X, Yew DT, Wu Y (2010) Amyloid-β induces caspase-dependent loss of PSD-95 and synaptophysin through NMDA receptors. J Alzheimers Dis 22, 541–56.

55. Ramírez-Hernández E, Sánchez-Maldonado C, Patricio-Martínez A, Limón ID (2023) Amyloid-β_25-35_ induces the morphological alteration of dendritic spines and decreases NR2B and PSD-95 expression in the hippocampus. Neurosci Lett 795, 137030.

56. Kornau HC, Schenker LT, Kennedy MB, Seeburg PH (1995) Domain interaction between NMDA receptor subunits and the postsynaptic density protein PSD-95. Science 269, 1737–40.

57. Niethammer M, Kim E, Sheng M (1996) Interaction between the C terminus of NMDA receptor subunits and multiple members of the PSD-95 family of membrane-associated guanylate kinases. J Neurosci 16, 2157–63.

58. Dore K, Carrico Z, Alfonso S, Marino M, Koymans K, Kessels HW, Malinow R (2021) PSD-95 protects synapses from β-amyloid. Cell Rep 35, 109194.

59. Pérez-Otaño I, Luján R, Tavalin SJ, Plomann M, Modregger J, Liu XB, Jones EG, Heinemann SF, Lo DC, Ehlers MD. Endocytosis and synaptic removal of NR3A-containing NMDA receptors by PAC-SIN1/syndapin1. Nat Neurosci 2006;9:611–21.

60. Yeung JHY, Walby JL, Palpagama TH, Turner C, Waldvogel HJ, Faull RLM, Kwakowsky A (2021) Glutamatergic receptor expression changes in the Alzheimer’s disease hippocampus and entorhinal cortex. Brain Pathol 31, e13005.

61. Köhr G (2006) NMDA receptor function: subunit composition versus spatial distribution. Cell Tissue Res 326, 439–46.

62. Vizi ES, Kisfali M, Lőrincz T (2013) Role of nonsynaptic GluN2B-containing NMDA receptors in excitotoxicity: evidence that fluoxetine selectively inhibits these receptors and may have neuroprotective effects. Brain Res Bull 93, 32–8.

63. Léveillé F, El Gaamouch F, Gouix E, Lecocq M, Lobner D, Nicole O, Buisson A (2008) Neuronal viability is controlled by a functional relation between synaptic and extrasynaptic NMDA receptors. FASEB J 22, 4258–71.

64. McQuate A, Barria A (2020) Rapid exchange of synaptic and extrasynaptic NMDA receptors in hippocampal CA1 neurons. J Neurophysiol 123, 1004–1014.

65. Lee HK, Min SS, Gallagher M, Kirkwood A (2005) NMDA receptor-independent long-term depression correlates with successful aging in rats. Nat Neurosci. 2005;8:1657–9.

66. Papadia S, Stevenson P, Hardingham NR, Bading H, Hardingham GE. Nuclear Ca^2+^ and the cAMP response element-binding protein family mediate a late phase of activity-dependent neuroprotection. J Neurosci 25, 4279–87.

67. Papouin T, Oliet SH (2014) Organization, control and function of extrasynaptic NMDA receptors. Philos Trans R Soc Lond B Biol Sci 369, 20130601.

73. 68. Müller MK, Jacobi E, Sakimura K, Malinow R, von Engelhardt J (2018) NMDA receptors mediate synaptic depression, but not spine loss in the dentate gyrus of adult amyloid Beta (Aβ) overexpressing mice. Acta Neuropathol Commun 23, 6:110.

69. Li Y, Sun W, Han S, Li J, Ding S, Wang W, Yin (2016) IGF-1-involved negative feedback of NR2B NMDA subunits protects cultured hippocampal neurons against NMDA-induced excitotoxicity. Mol Neurobiol 54, 684–96.

70. Ravikrishnan A, Gandhi PJ, Shelkar GP, Liu J, Pavuluri R, Dravid SM (2018) Region-specific expression of NMDA receptor GluN2C Subunit in parvalbumin-positive neurons and astrocytes: analysis of GluN2C expression using a novel reporter model. Neuroscience 380, 49–62.

71. Dingledine R, Borges K, Bowie D, Traynelis SF (1999) The glutamate receptor ion channels. Pharmacol Rev 51, 7–61.

77. 72. Kaindl AM, Degos V, Peineau S, Gouadon E, Chhor V, Loron G, Le Charpentier T, Josserand J, Ali C, Vivien D, Collingridge GL, Lombet A, Issa L, Rene F, Loeffler JP, Kavelaars A, Verney C, Mantz J, Gressens P (2012) Activation of microglial N-methyl-D-aspartate receptors triggers inflammation and neuronal cell death in the developing and mature brain. Ann Neurol, 72, 536–49.

73. Chou TH, Kang H, Simorowski N, Traynelis SF, Furukawa H (2022) Structural insights into assembly and function of GluN1-2C, GluN1-2A-2C, and GluN1-2D NMDARs. Mol Cell 82, 4548–4563.e4.

74. Alberdi E, Sánchez-Gómez MV, Cavaliere F, Pérez-Samartín A, Zugaza JL, Trullas R, Domercq M, Matute C (2010) Amyloid β oligomers induce Ca^2+^ dysregulation and neuronal death through activation of ionotropic glutamate receptors. Cell Calcium 47, 264–72.

75. Bolmont T, Haiss F, Eicke D, Radde R, Mathis CA, Klunk WE, Kohsaka S, Jucker M, Calhoun ME (2008) Dynamics of the microglial/amyloid interaction indicate a role in plaque maintenance. J Neurosci 28, 4283–92.

76. Eapen AV, Fernández-Fernández D, Georgiou J, Bortolotto ZA, Lightman S, Jane DE, Volianskis A, Collingridge GL (2021) Multiple roles of GluN2D-containing NMDA receptors in short-term potentiation and long-term potentiation in mouse hippocampal slices. Neuropharmacology 201, 108833.

77. Karimi Tari P, Parsons CG, Collingridge GL, Rammes G (2024) Memantine: Updating a rare success story in pro-cognitive therapeutics. Neuropharmacology 244, 109737.

78. Filali M, Lalonde R, Rivest S (2011) Subchronic memantine administration on spatial learning, exploratory activity, and nest-building in an APP/PS1 mouse model of Alzheimer’s disease. Neuropharmacology 60, 930–6.

79. Scholtzova H, Wadghiri YZ, Douadi M, Sigurdsson EM, Li YS, Quartermain D, Banerjee P, Wisniewski T (2008) Memantine leads to behavioral improvement and amyloid reduction in Alzheimer’s-disease-model transgenic mice shown as by micromagnetic resonance imaging. J Neurosci Res 86, 2784–91.

80. Li S, Jin M, Koeglsperger T, Shepardson NE, Shankar GM, Selkoe DJ (2011) Soluble Aβ oligomers inhibit long-term potentiation through a mechanism involving excessive activation of extrasynaptic NR2B-containing NMDA receptors. J Neurosci 31, 6627–38.

81. Minkeviciene R, Ihalainen J, Malm T, Matilainen O, Keksa-Goldsteine V, Goldsteins G, Iivonen H, Leguit N, Glennon J, Koistinaho J, Banerjee P, Tanila H (2008) Age-related decrease in stimulated glutamate release and vesicular glutamate transporters in APP/PS1 transgenic and wild-type mice. J Neurochem 105, 584–94.

87. 82. Puoliväli J, Wang J, Heikkinen T, Heikkilä M, Tapiola T, van Groen T, Tanila H (2002) Hippocampal Ab_42_ levels correlate with spatial memory deficit in APP and PS1 double transgenic mice. Neurobiol Dis 9, 339–47.

88. 83. Benilova I, Karran E, De Strooper B (2012) The toxic Aβ oligomer and Alzheimer’s disease: an emperor in need of clothes. Nat Neurosci 15, 349–357.

84. Goure WF, Krafft GA, Jerecic J, Hefti F (2014) Targeting the proper amyloid-beta neuronal toxins: a path forward for Alzheimer’s disease immunotherapeutics. Alzheimers Res Ther 6, 42.

85. Cheng IH, Scearce-Levie K, Legleiter J, Palop JJ, Gerstein H, Bien-Ly N, Puoliväli J, Lesné S, Ashe KH, Muchowski PJ, Mucke L (2007) Accelerating amyloid-β fibrillization reduces oligomer levels and functional deficits in Alzheimer disease mouse models. J Biol Chem 282, 23818–28.

86. Alsaad HA, DeKorver NW, Mao Z, Dravid SM, Arikkath J, Monaghan DT (2019) In the Telencephalon, GluN2C NMDA Receptor Subunit mRNA is Predominately Expressed in Glial Cells and GluN2D mRNA in Interneurons. Neurochem Res, 44, 61–77.

92. 87. Di Castro MA, Volterra A (2022) Astrocyte control of the entorhinal cortex-dentate gyrus circuit: Relevance to cognitive processing and impairment in pathology. Glia 70, 1536–1553.

88. Yoshiike Y, Kimura T, Yamashita S, Furudate H, Mizoroki T, Murayama M, Takashima A (2008) GABA_A_ receptor-mediated acceleration of aging-associated memory decline in APP/PS1 mice and its pharmacological treatment by picrotoxin. PLoS One 3, e3029.

89. Ge S, Yang CH, Hsu KS, Ming GL, Song H (2007) A critical period for enhanced synaptic plasticity in newly generated neurons of the adult brain. Neuron 54, 559–66.

95. 90. Ji Q, Yang Y, Xiong Y, Zhang YJ, Jiang J, Zhou LP, Du XH, Wang CX, Zhu ZR (2023) Blockade of adenosine A2A receptors reverses early spatial memory defects in the APP/PS1 mouse model of Alzheimer’s disease by promoting synaptic plasticity of adult-born granule cells. Alzheimers Res Ther 15, 187.

